# Inhibition of muscarinic receptor signaling protects human enteric inhibitory neurons against platin chemotherapy toxicity

**DOI:** 10.1101/2023.03.08.531806

**Authors:** Mikayla N Richter, Sina Farahvashi, Ryan M Samuel, Homa Majd, Angeline K Chemel, Jonathan T Ramirez, Alireza Majd, Megan D Scantlen, Nicholas Elder, Andrius Cesiulis, Kristle Garcia, Tanvi Joshi, Matthew G Keefe, Bardia Samiakalantari, Elena M Turkalj, Johnny Yu, Abolfazl Arab, Keyi Yin, Bruce Culbertson, Bianca Vora, Chenling Xiong, Michael G Kattah, Roshanak Irannejad, Deanna L Kroetz, Tomasz J Nowakowski, Hani Goodarzi, Faranak Fattahi

**Affiliations:** Department of Cellular and Molecular Pharmacology, University of California, San Francisco, San Francisco, California, USA; Eli and Edythe Broad Center of Regeneration Medicine and Stem Cell Research, University of California, San Francisco, California, USA; Department of Biochemistry and Biophysics, University of California, San Francisco, San Francisco, California, USA; Department of Anatomy, University of California, San Francisco, San Francisco, California, USA; Department of Psychiatry and Behavioral Sciences, University of California, San Francisco, California, USA; Department of Bioengineering and Therapeutic Sciences, University of California, San Francisco, San Francisco, California, USA; Department of Medicine, University of California, San Francisco, San Francisco, California, USA; Department of Neurological Surgery, University of California, San Francisco, San Francisco, California, USA; Department of Urology, University of California, San Francisco, San Francisco, California, USA; Program in Craniofacial Biology, University of California, San Francisco, California, USA

## Abstract

GI toxicity is a common dose-limiting adverse effect of platin chemotherapy treatment. Up to 50% of cancer survivors continue to experience symptoms of chronic constipation or diarrhea induced by their chemotherapy for many years after their treatment. This drug toxicity is largely attributed to damage to enteric neurons that innervate the GI tract and control GI motility. The mechanisms responsible for platin-induced enteric neurotoxicity and potential preventative strategies have remained unknown. Here, we use human pluripotent stem cell derived enteric neurons to establish a new model system capable of uncovering the mechanism of platin-induced enteric neuropathy. Utilizing this scalable system, we performed a high throughput screen and identified drug candidates and pathways involved in the disease. Our analyses revealed that excitotoxicity through muscarinic cholinergic signaling is a key driver of platin-induced enteric neuropathy. Using single nuclei transcriptomics and functional assays, we discovered that this disease mechanism leads to increased susceptibility of specific neuronal subtypes, including inhibitory nitrergic neurons, to platins. Histological assessment of the enteric nervous system in platin-treated patients confirmed the selective loss of nitrergic neurons. Finally, we demonstrated that pharmacological and genetic inhibition of muscarinic cholinergic signaling is sufficient to rescue enteric neurons from platin excitotoxicity *in vitro* and can prevent platin-induced constipation and degeneration of nitrergic neurons in mice. These studies define the mechanisms of platin-induced enteric neuropathy and serve as a framework for uncovering cell type-specific manifestations of cellular stress underlying numerous intractable peripheral neuropathies.

## Introduction

Platin chemotherapies, including cisplatin, carboplatin, and oxaliplatin, have been approved and used widely to treat cancer for decades. They inhibit cancer progression by binding DNA, forming DNA adducts that induce DNA strand breaks, which ultimately lead to cell death if the damage is overwhelming^1^. This mechanism of action is highly effective at treating a variety of cancers. For example, since the introduction of cisplatin in 1978, the prognosis of testicular, ovarian, and bladder cancers has improved tremendously (Seer Cancer Statistics Review 1975-2015). Furthermore, platins are prescribed for highly prevalent cancers, including colorectal, lung, and bladder cancers, making up about 50% of all cancer patients^2^. However, despite their clinical efficacy and applicability, a major adverse effect that limits the utility of platins is gastrointestinal (GI) dysmotility, resulting from platin-induced damage to the enteric nervous system (ENS)^3–6^.

The ENS is the division of the peripheral nervous system embedded in the GI tract that regulates GI functions, including motility, secretion, and immune response^7^. It contains a diverse network of neurons that express specific neurotransmitters and neuropeptides corresponding with distinct functional roles, including excitatory and inhibitory activity^8^. Responding to local stimuli in the lumen, the ENS neuronal network coordinates the contraction and relaxation of smooth muscle fibers to orchestrate complex patterns of motility in different parts of the GI tract. Disruptions to one or more components of the circuit can lead to motility disorders including achalasia, gastroparesis, intestinal pseudo-obstruction, chronic constipation, or chronic diarrhea^9–11^.

Platin-induced GI toxicities replicate many symptomatic aspects of GI motility disorders, including, most commonly, platin-induced constipation or diarrhea. For example, oxaliplatin causes diarrhea in over 20% of patients and constipation in over 60% of patients, as reported during clinical trials^5,6^. Importantly, 49% of patients report continuing symptoms post-treatment, suggesting damage to a non-regenerative tissue, such as neurons^12^. This hypothesis has been supported by recent reports of platin-induced damage to the ENS in mice, including increases in mitochondrial stress, reactive oxygen species levels, and disrupted mitochondrial membrane integrity which coincide with a decrease in the total number of enteric neurons^13–18^. These studies also report no significant damage to the muscle or mucosal tissue along the GI tract, which aligns with the low incidence of mucositis reported in humans^4–6^. Furthermore, there is evidence that platin-induced enteric neuropathy may be cell type-specific. For example, platins affect the proportion of enteric neuron subtypes in the ENS and the functional properties of nitrergic neurons, which are inhibitory enteric neurons^13,14,16,17,19^. Notably, nitrergic neuron-mediated electrophysiological properties and motility phenotypes are disrupted following oxaliplatin treatment, indicating that inhibitory motor neuron functions may be specifically affected by platins^14^. Ultimately, platins cause colonic motility to become increasingly ineffective in mice, manifesting in the clinically relevant phenotype of constipation^13,14,17,18^.

These studies have solidified the link between the clinical manifestations of platin-induced GI dysmotility and cellular phenotypes of enteric neuropathy. Unfortunately, despite having a robust mouse model, the molecular pathways underlying the toxicity mechanism remain unidentified and no druggable targets have been nominated. Furthermore, it remains unclear why certain enteric neurons may be affected more than others. This is largely because of the technical challenges and translational limitations of studying platin-induced enteric neuropathy in animals. For example, enteric neurons reside solely within the GI tract, where they make up about 1% of the tissue making them inaccessible and difficult to study at large scale^20^. Furthermore, mouse models and primary tissue samples are not scalable and therefore incompatible with high throughput methodologies such as unbiased pharmacologic screening that enable the discovery of disease mechanisms and drug targets.

Here, we utilized our established method of deriving enteric neurons from human pluripotent stem cells (hPSCs)^21–23^ to build a robust *in vitro* model of platin-induced enteric neuropathy and uncover the mechanism of toxicity. Using this system, we determined the cellular and molecular hallmarks of platin-induced enteric neuropathy through comprehensive phenotypic and transcriptomic profiling. Leveraging this model, we performed a high throughput screen, defined the toxicity mechanism and identified drug candidates and potential therapeutic targets. The results revealed the excitotoxicity signaling network responsible for cell type specific susceptibility of nitrergic neurons to platins. Finally, we demonstrated that blocking this disease pathway using our drug candidate can prevent platin-induced enteric neuropathy in mice.

## Results

### Enteric neurons are directly susceptible to platin chemotherapies

Despite platin chemotherapy drugs being approved and in use for decades, their adverse effects on GI functions have only been documented in clinical trials. To evaluate the prevalence of these adverse effects in the real world, we leveraged electronic medical record data from the UCSF Research Data Browser to investigate the propensity of a patient to develop chemotherapy-induced constipation after undergoing oxaliplatin or carboplatin treatment. For both oxaliplatin and carboplatin, we followed a similar analysis workflow (Figure S1A). We identified and extracted patients diagnosed with colorectal cancer (for the oxaliplatin analysis) or ovarian cancer (for the carboplatin analysis). To limit confounding the data, in both analyses, we excluded all patients who were diagnosed with constipation prior to their cancer diagnosis. The resulting patients in the platin chemotherapy treated and untreated groups were then matched based on age, sex, and race/ethnicity for each analysis. This ultimately resulted in sample sizes of 215 patients with colorectal cancer (46 patients prescribed oxaliplatin, 169 patients not prescribed oxaliplatin) and 4087 patients with ovarian cancer (438 patients prescribed carboplatin, 3649 patients not prescribed carboplatin), which were used for our analysis (Supplementary Tables 1 and 2). The patients were then subset into four groups based on whether they were diagnosed with constipation. Patients prescribed oxaliplatin are 3.4 times more likely to develop constipation compared to patients not prescribed oxaliplatin with a p-value of < 0.001 (Figure S1B and Supplementary Table 3). Similarly, patients prescribed carboplatin are 8.2 times more likely to be diagnosed with constipation (compared to patients not prescribed carboplatin) with a p-value of < 0.001 (Figure S1B and Supplementary Table 3). These data highlight the detrimental effect of this chemotherapy drug class on GI motility. Furthermore, in our analysis, 57% of patients prescribed oxaliplatin and 49% of patients prescribed carboplatin were diagnosed with constipation, which does coincide with percentages reported in some clinical trials^4,5^. However, our calculations are much higher than the prevalence of constipation reported in the package inserts for oxaliplatin and carboplatin, 31% and 6%, respectively, highlighting potential inconsistencies between the reported drug adverse effects and the actual prevalence of the adverse effect in a large, more diverse, real world patient population.

Animal studies have established the role of ENS damage in platin-induced GI dysmotility^13–18^, but it remains unclear if this toxicity is due to a potentially higher exposure of the ENS tissue during drug delivery or an inherent susceptibility of enteric neurons to platins. We sought to address this possibility by exposing different peripheral neurons to identical platin treatment conditions and compare their response. Due to the limitations associated with isolation and maintenance of human neurons from primary tissue, we used our previously established hPSC differentiation system to generate human enteric neurons^21–23^. Following approximately 40 days of differentiation, these hPSC-derived enteric neurons express a diverse array of neurotransmitter subtype markers, indicative of their successful recapitulation of the human ENS (Figure S1C). Similarly, we utilized an established protocol for generating cranial neural crest-derived peripheral neuron lineages from hPSCs^24^. Following approximately 40 days of differentiation, this protocol can successfully generate a mixture of ASCL1+ and TH+ sympathetic neurons, BRN3A+ sensory neurons, and CHAT+ parasympathetic neurons, making it an ideal peripheral neuron control group to compare with enteric neurons (Figure S1C).

Leveraging these *in vitro* models, we aimed to characterize the direct effects platins have on enteric neurons as compared to control peripheral neurons (Figure 1A). Platin-induced cytotoxicity was measured in both neuronal populations by measuring the activity of lactate dehydrogenase (LDH) released by dead cells in the cell culture supernatant. For each platin, the three day treatment resulted in the enteric neuron population having a significantly higher cytotoxicity response relative to the peripheral neuron control group, suggesting that enteric neurons are more vulnerable to platin toxicity relative to other peripheral neurons (Figure 1B). No differences in cytotoxicity were observed when the two neuron populations were treated with the mitochondrial toxin carbonyl cyanide m-chlorophenyl hydrazone (CCCP), indicating that the differential sensitivity to platins is not due to a higher general susceptibility to cellular stress.

**Figure 1:**
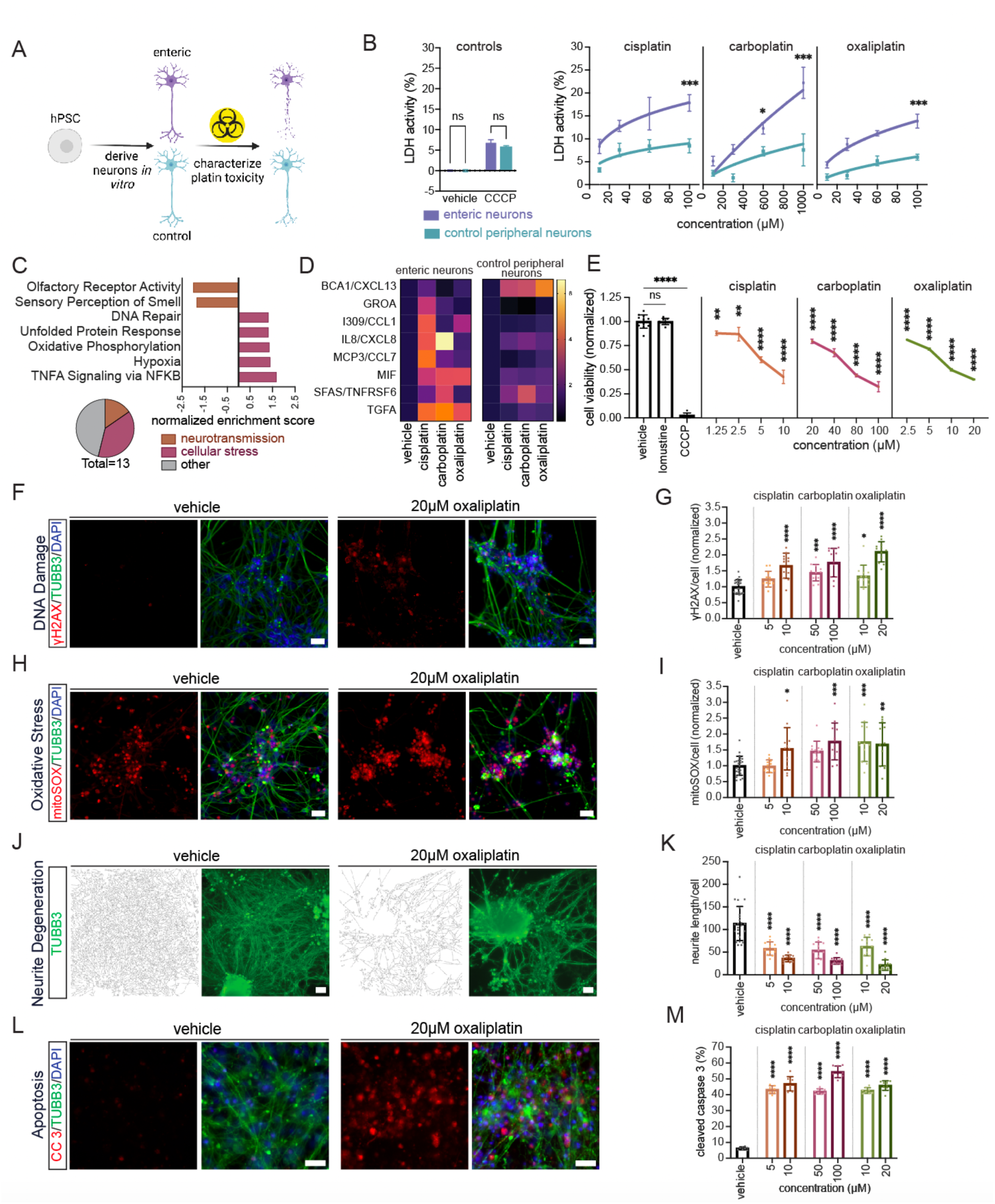
Enteric neurons are sensitively vulnerable to platin chemotherapies. **A)** Schematic illustration of *in vitro* characterization of platin toxicity in enteric and control peripheral neurons. **B)** Lactate dehydrogenase cytotoxicity analysis comparing the response of hPSC-derived enteric neurons to hPSC-derived control peripheral neurons treated with vehicle (medium containing 10% water), 100 μM CCCP, and increasing platin concentrations. All values are normalized by subtracting the vehicle condition from each sample. Data are represented as mean ± SEM. **C)** Summary of gene sets significantly enriched in the log_2_ fold-change gene expression list comparing 50 μM oxaliplatin-treated enteric neurons to 50 μM oxaliplatin-treated control peripheral neurons. The bar graph highlights neurotransmission and cellular stress related pathways and the pie chart shows the proportion of significantly enriched gene sets relating to neurotransmission or cellular stress. **D)** Heatmap of the cytokines significantly increased in the cell culture supernatant of enteric and control peripheral neurons treated with 20 µM cisplatin, 200 µM carboplatin, 40 µM oxaliplatin, or medium containing 4% water (vehicle). Each cytokine is normalized to its vehicle cytokine levels. **E)** Cell viability analysis of hPSC-derived enteric neurons exposed to vehicle (medium containing 2% water), 20 μM lomustine, 10 μM CCCP, and increasing platin concentrations. All values are normalized to vehicle. Data are represented as mean ± SEM. **F, G**) γH2AX staining and quantification in platin-treated hPSC-derived enteric neurons. All values are normalized to vehicle (medium containing 2% water). Scale bar: 50 μm. Data are represented as mean ± SD. **H, I**) mitoSOX staining and quantification in platin-treated hPSC-derived enteric neurons. All values are normalized to vehicle (medium containing 2% water). Scale bar: 50 μm. Data are represented as mean ± SD. **J, K**) TUBB3 staining, TUBB3 neurite traces, and quantification in platin-treated hPSC-derived enteric neurons. Vehicle condition contains water diluted to 2% in medium. Scale bar: 50 μm. Data are represented as mean ± SD. **L, M**) Cleaved caspase 3 staining and quantification in platin-treated hPSC-derived enteric neurons. Vehicle condition contains water diluted to 2% in medium. Scale bar: 50 μm. Data are represented as mean ± SD. *p value < 0.05, ** p value < 0.01, *** p value < 0.001, **** p value < 0.0001, ns: not significant.

To further characterize this cell-type specific response to platins, we performed bulk transcriptomic profiling of the enteric neuron and the control peripheral neuron populations. Read counts of oxaliplatin treated enteric neurons were compared to the oxaliplatin treated control peripheral neurons to calculate the log_2_ fold-change in gene expression. Then pre-ranked gene set enrichment analysis (GSEA) was performed to identify gene sets significantly changed between the two populations based on a p-value cutoff of < 0.05 and a false discovery rate of < 0.25. Multiple gene sets related to cellular stress were positively enriched in the enteric neurons relative to the control peripheral neurons, further suggesting enteric neurons have an increased susceptibility to platin toxicity (Figure 1C). These cellular stress gene sets included oxidative phosphorylation, which may reflect the phenotype of increased oxaliplatin-induced mitochondrial oxidative stress observed *in vivo* ^13,14^. The DNA repair pathway likely reflects the cellular response to oxaliplatin’s mechanism of action, DNA damage^1^. Other cellular stress pathways such as unfolded protein response, hypoxia, and TNF signaling via NF-κB have never been characterized in the context of platin-induced enteric neuropathy, and may reflect other neurotoxic pathways ongoing during drug exposure. Additionally, two neurotransmission related pathways were negatively enriched in the enteric neurons relative to the control peripheral neurons: olfactory receptor activity and sensory perception of smell. This suggests that platins may impact neuronal signaling in enteric neurons more than other peripheral neurons and this may be another contributing factor to platin-induced enteric neuropathy.

To identify biological pathways transcriptionally dysregulated by platins in hPSC-derived enteric neurons, we performed pre-ranked GSEA on the log_2_ fold-change in gene expression between oxaliplatin treated enteric neurons and vehicle treated enteric neurons. Many gene sets related to neuronal signaling pathways were significantly negatively enriched, indicating that aberrant neurotransmission may be a key pathway underlying platin-induced enteric neuropathy (Figure S1D). Furthermore, many cellular stress gene sets reflecting neuroinflammatory, oxidative stress, and DNA damage pathways were again revealed in this analysis (Figure S1D). As a control, we also performed this analysis on the log_2_ fold-change in gene expression between oxaliplatin treated control peripheral neurons and vehicle treated control peripheral neurons. None of the significantly enriched gene sets reflect neurotransmission or cellular stress related pathways, again suggesting that they are less susceptible to platin toxicity (Figure S1E).

The significant enrichment of TNF signaling via NF-κB in enteric neurons relative to other peripheral neurons indicates that neuroinflammation may have a role in potentiating platin-induced enteric neuropathy. NF-κB is a master regulator of neuroinflammation that ultimately can lead to the release of cytokines and chemokines from the cell to activate the appropriate immune cells to clear the inflammation and tissue damage^25^. To experimentally test if platin-induced neuroinflammation is increased in enteric neurons relative to other peripheral neurons, we performed a cytokine array detecting the levels of 76 neuroinflammatory cytokines in each cell population’s culture supernatant after three days exposure to the platins (Supplementary Table 4). The median fluorescence intensities for each cytokine were averaged and normalized to each population’s vehicle treatment condition. Enteric neurons significantly increased the release of seven inflammatory cytokines in response to at least one platin, including C-X-C motif chemokine ligand 1 (GROA), chemokine C-C motif ligand 1 (I309/CCL1), interleukin-8 (IL8/CXCL8), chemokine C-C motif ligand 7 (MCP3/CCL7), macrophage migration inhibitory factor (MIF), tumor necrosis factor receptor superfamily member 6 (SFAS/TNFRSF6), and transforming growth factor alpha (TGFA), whereas the control peripheral neuron population, on the other hand, only had two cytokines that were significantly increased: chemokine C-X-C motif ligand 13 (BCA1/CXCL13) and SFAS/TNFRSF6 (Figure 1D). These results confirm the transcriptomic-based prediction of enteric neurons experiencing an increased neuroinflammatory response to platins relative to other peripheral neurons (Figure 1C). Furthermore, SFAS/TNFRSF6 is the only cytokine significantly increased in both populations (Figure 1D). This indicates that the enteric neuron inflammatory response to platins is largely non-overlapping with other peripheral neurons and reflects a cell-type specific drug response. This highlights the importance of using a model system that best reflects the distinct molecular state of the relevant cell type.

The evidence that enteric neurons are significantly more sensitive to platin chemotherapies as compared to other divisions of the peripheral nervous system prompted us to systematically evaluate the neurotoxic phenotypes platin chemotherapies directly induce in hPSC-derived enteric neurons. To better characterize the effect of platins on enteric neuron viability, we treated enteric neurons with a range of platin chemotherapy concentrations for three days. We observed dose-dependent decreases in cell viability, measured via ATP levels, with LC_50_ values of 4.7 μM, 69.3 μM, and 7.1 μM and for cisplatin, carboplatin, and oxaliplatin, respectively (Figure 1E). To determine if this reduction in enteric neuron viability is a platin-specific mechanism or not, we also exposed enteric neurons to lomustine, a chemotherapy drug with a similar mechanism of action to platins, but with low incidence of GI toxicity reported clinically^26^. We observed no significant effects on enteric neuron viability when hPSC-derived enteric neurons were treated with lomustine for three days (Figure 1E). This data suggests that the enteric neurons are directly sensitive to platin chemotherapies via a platin chemotherapy-specific toxicity mechanism.

To validate that these *in vitro* treatment conditions reflect exposures a patient might experience clinically, we calculated patient exposure by dividing the recommended patient dose by each platin’s clearance. Performing this calculation for each indication the platin is approved for, we generated a range of expected patient exposures for each platin chemotherapy (Figure S1F). *In vitro* exposures were calculated by multiplying the concentrations used *in vitro* by the duration of the experiment – 72 hours. These *in vitro* exposures overlap with the patient exposures, further indicating that the *in vitro* experiments reflect biologically and clinically relevant conditions (Figure S1F).

To experimentally validate the oxaliplatin-induced cellular stress response predicted from our transcriptomic profiling, we performed relevant phenotypic assays in our hPSC-derived enteric neuron cultures. Significant positive enrichment of the DNA repair and p53 pathway gene sets suggests platins directly induce DNA damage in enteric neurons (Figure 1C and S1D). To measure levels of DNA damage, we performed immunostaining to detect γH2AX, a histone post-translational modification that occurs when a DNA double strand break is detected^27^. All three platins significantly increased γH2AX levels in hPSC-derived enteric neurons, confirming that platins indeed cause DNA damage in enteric neurons (Figure 1F-G). Positive enrichment of the oxidative phosphorylation and mitochondrial large ribosomal subunit gene sets suggests platins may directly induce mitochondrial oxidative stress in enteric neurons (Figure 1C and S1D). Increased mitochondrial reactive oxygen species and mitochondrial membrane permeability has been observed in mouse models of oxaliplatin-induced enteric neuropathy^13,14^. However, in these studies it remained unclear if this phenotype was due to direct or indirect effects of platins on enteric neurons or changes in the vascular tissue that supply oxygen to the ENS tissue. Our transcriptomic profiling on hPSC-derived enteric neurons *in vitro* suggests that this toxicity hallmark is conserved in human cells and is due to a direct effect of platins on enteric neurons. To test this hypothesis, we utilized the mitochondrial superoxide indicator, mitoSOX, to detect oxidative stress in platin-treated enteric neurons. Indeed, mitochondrial superoxide levels were significantly increased following platin exposure in hPSC-derived enteric neurons, confirming the results of our transcriptomic profiling and supporting the role of increased reactive oxygen species in potentiating platin-induced enteric neuropathy (Figure 1H-I). Given that neurodegeneration is a common hallmark of neuropathy and has been observed in other forms of chemotherapy-induced neurotoxicity^28–30^, we aimed to identify if platins induce neurodegeneration in enteric neurons. We characterized platin-induced neurodegeneration by quantifying the total neurite length per cell, measured by TUBB3 immunostaining and DAPI staining, respectively. Through this method, we observed that each platin indeed causes reduces the neurite networks in hPSC-derived enteric neurons, indicating that platins cause enteric neurons to degenerate (Figure 1J-K). Lastly, the positive enrichment of the apoptosis and p53 pathway gene sets suggests platins induce enteric neuron cell death via apoptosis (Figure S1D). To measure apoptosis induction, we detected levels of cleaved caspase 3, a post-translational modification that occurs to initiate apoptosis^31^. At each concentration tested, all three platins significantly increased the percentage of neurons positive for cleaved caspase 3, indicating the initiation of apoptosis (Figure 1L-M). Thus, platins directly induce DNA damage and oxidative stress in hPSC-derived enteric neurons, leading to neurite degeneration and programmed cell death.

Altogether, these data indicate that enteric neurons are directly sensitive to platin chemotherapies and suggests that platin-induced enteric neuropathy may be largely explained by the direct interaction between platins and enteric neurons, as all phenotypes were induced *in vitro*, in the absence of the *in vivo* tissue micro and macro environments. Therefore, methods that block the direct effect of platins on enteric neurons may be highly effective at preserving this vulnerable cell type.

### Phenotypic screen identifies platin-induced excitotoxicity mechanism

Leveraging our scalable model of platin-induced enteric neuropathy, we carried out a high throughput phenotypic drug screen followed by drug target enrichment analysis to uncover the molecular mechanism underlying platin-induced enteric neuropathy (Figure 2A). Searching for modulators of oxaliplatin-induced cell death, we screened the Selleckchem small molecule library of 1,443 FDA-approved drugs in both iPSC and ESC-derived enteric neurons (Figure 2B). Cell viability was quantified by detecting dead cells via DAPI staining prior to fixation and all cells via propidium iodide staining post fixation. From the screen, we confirmed the effect of three compounds with high positive cell viability z-scores (mefloquine, mitotane, and oxybutynin). All were able to significantly reduce oxaliplatin-induced cell death, measured via ATP levels as an orthogonal cell viability detection method, suggesting they can block the toxicity mechanism and increase cell viability when dosed in combination with oxaliplatin (Figure 2C).

**Figure 2:**
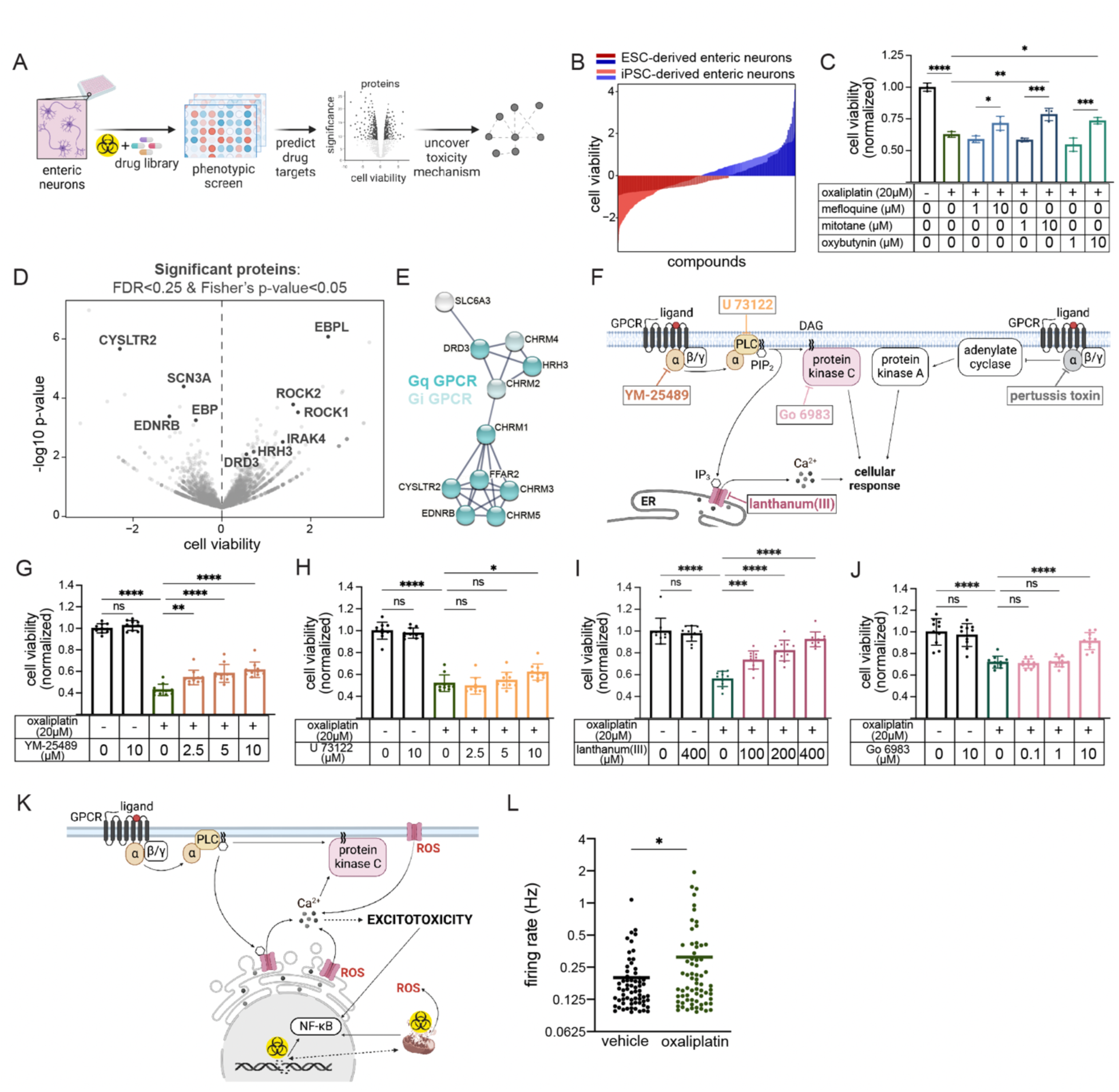
High throughput screen uncovers mechanism of platin-induced excitotoxicity in enteric neurons. **A)** Schematic illustration of our drug and drug target discovery pipeline. High throughput screening followed by SEA-STRING analysis identifies drugs and drug targets that affect hPSC-derived enteric neuron viability during oxaliplatin treatment. **B)** High throughput screening of Selleckchem FDA approved drug library identifies drugs that increase and decrease hPSC-derived enteric neuron viability during oxaliplatin treatment. **C)** Dose response of the hit compounds on hPSC-derived enteric neuron viability when dosed in combination with oxaliplatin. All values are normalized to vehicle (medium containing 2% water). Data are represented as mean ± SD. **D)** Volcano plot showing the average cell viability z-score versus the −log_10_ p-value of the genes targeted in the screen. The genes that pass our statistical thresholds are marked in black (combined z-score FDR < 0.25 and Fisher’s p-value < 0.05). **E)** STRING protein-protein interaction network formed from our gene list of predicted drug targets. Network is primarily composed of GPCRs with proteins of the G_q_ class shown in dark cyan and proteins of the G_i_ class shown in light cyan. The minimum required interaction score was set to 0.7, reflecting high confidence interactions. **F)** Schematic illustration of our strategy to use tool compounds to interrogate the role of the G_q_-mediated inositol phospholipid signaling pathway and the G_i_-mediated cyclic adenosine monophosphate dependent pathway in potentiating platin-induced enteric neuropathy. **G-J**) Dose response of the tool compounds on hPSC-derived enteric neuron viability when dosed in combination with oxaliplatin. All values are normalized to vehicle (medium containing 2% water). (**G**) YM-25489 a G_q_ inhibitor, (**H**) U 73122, a phospholipase C inhibitor, (**I**) lanthanum (III), a calcium channel blocker potent at IP3 receptors, (**J**) Go 6983, a protein kinase C inhibitor. Data are represented as mean ± SD. **K)** Schematic illustration of our platin-induced excitotoxicity mechanism. **L)** Quantification of microelectrode array analysis of 10 μM oxaliplatin and vehicle (medium containing 1% water)-treated hPSC-derived enteric neurons. Data are represented as mean. *p value < 0.05, ** p value < 0.01, *** p value < 0.001, **** p value < 0.0001, ns: not significant.

To evaluate the clinical relevance of these compounds as therapies to protect enteric neurons during chemotherapy treatment, we evaluated them in two colorectal cancer cell lines (SW480 and WiDr), assessing whether any of them alter the antineoplastic efficacy of platin chemotherapies. In both colorectal cancer lines, mefloquine increased cell death when dosed in combination with the platins (Figure S2A, S2B, S2C). This result either suggests that mefloquine has antineoplastic activity itself or that it is generally toxic at the concentrations tested. Mitotane, on the other hand, consistently decreased cancer cell death when dosed in combination with the platins, suggesting that it likely protects enteric neurons by blocking the antineoplastic mechanism of the platins and therefore is an undesirable therapeutic candidate to counteract enteric neurotoxicity in cancer patients (Figure S2A, S2B, S2C). Oxybutynin exhibited no effect on the antineoplastic efficacy of any of the platins in either of the colorectal cancer cell lines, except for one exception. Oxybutynin did significantly decrease cell death when dosed at 4 µM in the WiDr line in combination with oxaliplatin, but this result was not replicated at the 1, 7 or 10 µM concentrations, so was likely driven by outliers (Figure S2C). Given that oxybutynin protects enteric neurons from platin-induced cell death without affecting cancer killing, its protective mechanism is likely DNA damage independent and reflects a parallel mechanism capable of preserving enteric neurons. Thus, oxybutynin is the compound from the drug screen that is most likely to be compatible with clinical dosing in cancer patients for preventing platin-induced GI neurotoxicity.

To identify the proteins that modulate oxaliplatin-induced enteric neuron viability, we utilized our previously published drug-protein interaction analysis pipeline to predict drug targets enriched in the compounds that either increase or decrease oxaliplatin-treated enteric neuron viability in both of our drug screen datasets^32^. Via this method, we identified 10 likely target proteins that met the significance criteria of a false discovery rate < 0.25 based on the combined z-score analysis and a Fisher’s p-value < 0.05 (Figure 2D and Supplementary Table 5). In addition to this unbiased approach, we also predicted the drug-protein interactions of our three validated drugs (mefloquine, mitotane, and oxybutynin), by inputting their simplified molecular-input line-entry system (SMILES) into the similarity ensemble approach (SEA) computational tool^33^. We filtered out the significant drug-protein interactions by selecting human proteins and predicted interaction p-values < 0.05, yielding 11 proteins predicted to be targeted by the validated hits (Supplementary Table 6). Through these unbiased and biased prediction methods, we identified a total of 21 potential targets.

To explore if the drug target proteins are part of shared pathway, we conducted a protein-protein interaction network analysis using the STRING physical interaction database^34^ to uncover a network of proteins with a high confidence interaction score. This approach revealed a core network of G protein-coupled receptors (GPCRs) of the G_q_ class shown in dark cyan and GPCRs of the G_i_ class shown in light cyan in Figure 2E, suggesting that GPCR signaling may potentiate platin toxicity in enteric neurons.

The convergence of the GPCR protein-protein interaction network with the neurotransmission pre-ranked GSEA results, led us to hypothesize that platin-induced enteric neuropathy may be reduced by antagonizing neurotransmitter-mediated GPCR signaling. To test this hypothesis, we employed five chemical tools that either inhibit or antagonize proteins along the G_q_-mediated inositol phospholipid signaling pathway and the G_i_-mediated cyclic adenosine monophosphate dependent pathway and evaluated their effects on oxaliplatin-induced cell viability (Figure 2F). To interrogate the role of the G_q_ protein alpha subunit, we utilized YM-25489, a G_q_ inhibitor, which when dosed in combination with oxaliplatin at 2.5, 5, and 10 µM was able to significantly increase cell viability, implicating the G_q_-mediated inositol phospholipid signaling pathway in platin toxicity in enteric neurons (Figure 2G). To interrogate the role of the G_i_ protein alpha subunit, we utilized pertussis toxin, a G_i_ inhibitor, which when dosed up to 0.5 μg/mL in combination with oxaliplatin was unable to significantly increase cell viability, suggesting the G_i_-mediated cyclic adenosine monophosphate dependent pathway is not a key driver of the platin toxicity mechanism (Figure S2D). Thus, we hypothesized that G_q_-mediated inositol phospholipid signaling is the primary contributor to the platin-induced enteric neuropathy mechanism and tested this hypothesis by inhibiting or antagonizing downstream effectors of the G_q_ protein. U 73122, a phospholipase C (PLC) inhibitor, was able to significantly increase enteric neuron viability when dosed at 10 μM in combination with oxaliplatin, indicating that PLC activated by G_q_ proteins is involved in the toxicity mechanism (Figure 2H). Active PLCs cleave phosphatidylinositol bisphosphate (PIP2), increasing inositol triphosphate (IP3) concentrations in the cytosol, which can bind and open IP3 calcium channels in the endoplasmic reticulum^35^. To explore the role of IP3 receptors in platin-induced enteric neuropathy, we utilized lanthanum (III), a calcium channel blocker potent at IP3 receptors, which also was able to significantly increase cell viability when dosed at 100, 200 and 400 μM in oxaliplatin-treated enteric neurons, suggesting that calcium is an important second messenger in the toxicity mechanism (Figure 2I). Lastly, when cleaving PIP2, PLCs also increase membrane-bound diacylglycerol (DAG) levels, which can activate protein kinase C (PKC) signaling^35^. To evaluate the role of PKC signaling in the platin-induced enteric neuropathy mechanism, we utilized Go 6983, a PKC inhibitor, which also protected enteric neuron cell viability when dosed in combination with oxaliplatin at 10 μM (Figure 2J). Together, these data suggest that platin-induced enteric neuropathy is potentiated by aberrant G_q_-mediated signaling.

Combining our identified hallmarks of platin-induced enteric neuropathy with the components of the inositol phospholipid signaling pathway, we formulated a potential mechanism of enteric neuron excitotoxicity triggered by platin-induced cellular stress (Figure 2K). The mechanism begins with platin-induced DNA damage and increased reactive oxygen species, which we observed *in vitro* and others have observed *in vivo* (Figure 1G-J)^13,14^. Reactive oxygen species can interact with calcium channels of the endoplasmic reticulum and extracellular membrane, permitting the depolarization of the cell, which, in neurons, leads to action potentials^36^. For enteric neurons with GPCRs of the G_q_ class, the G_q_-mediated inositol phospholipid signaling pathway leads to further depolarization of the cell, causing these neurons to enter an excitotoxicity positive feedback loop, which can lead to aberrant PKC activity and increased NF-κB-mediated neuroinflammatory activity.

To evaluate this hypothesis, we determined if platins increase the electrical activity of enteric neurons by performing a microelectrode array experiment measuring the mean firing rate in oxaliplatin-treated cultures. After a three hour incubation, oxaliplatin significantly increased the firing rate in treated neurons, as compared to vehicle-treated controls, suggesting enteric neurons indeed become hyperexcitable following oxaliplatin exposure (Figure 2L). Altogether, these data support our hypothesis that platin-induced enteric neuropathy is driven by calcium-mediated excitotoxicity.

### Muscarinic cholinergic signaling promotes platin-induced enteric neurotoxicity

Oxybutynin, one of the protective and most clinically relevant compounds from our drug screen (Figure 2C, S2A-C) is a potent muscarinic cholinergic receptor antagonist commonly prescribed for overactive bladder^37^. Acetylcholine is a key neurotransmitter in the ENS, mediating the activity of intrinsic circuits important for proper GI motility^7^. Notably, oxybutynin can decrease GI motility in humans, suggesting that its anticholinergic activity affects the ENS^37^. Three out of the five muscarinic cholinergic receptors are GPCRs of the G_q_ class, muscarinic cholinergic receptor 1 (CHRM1), muscarinic cholinergic receptor 3 (CHRM3), and muscarinic cholinergic receptor 5 (CHRM5), which led us to hypothesize that oxybutynin prevents platin-induced excitotoxicity by antagonizing G_q_ muscarinic cholinergic receptors. To test this hypothesis, we used both pharmacologic and genetic tools to determine oxybutynin’s therapeutic target and characterize its ability to prevent platin-induced excitotoxicity (Figure 3A).

**Figure 3:**
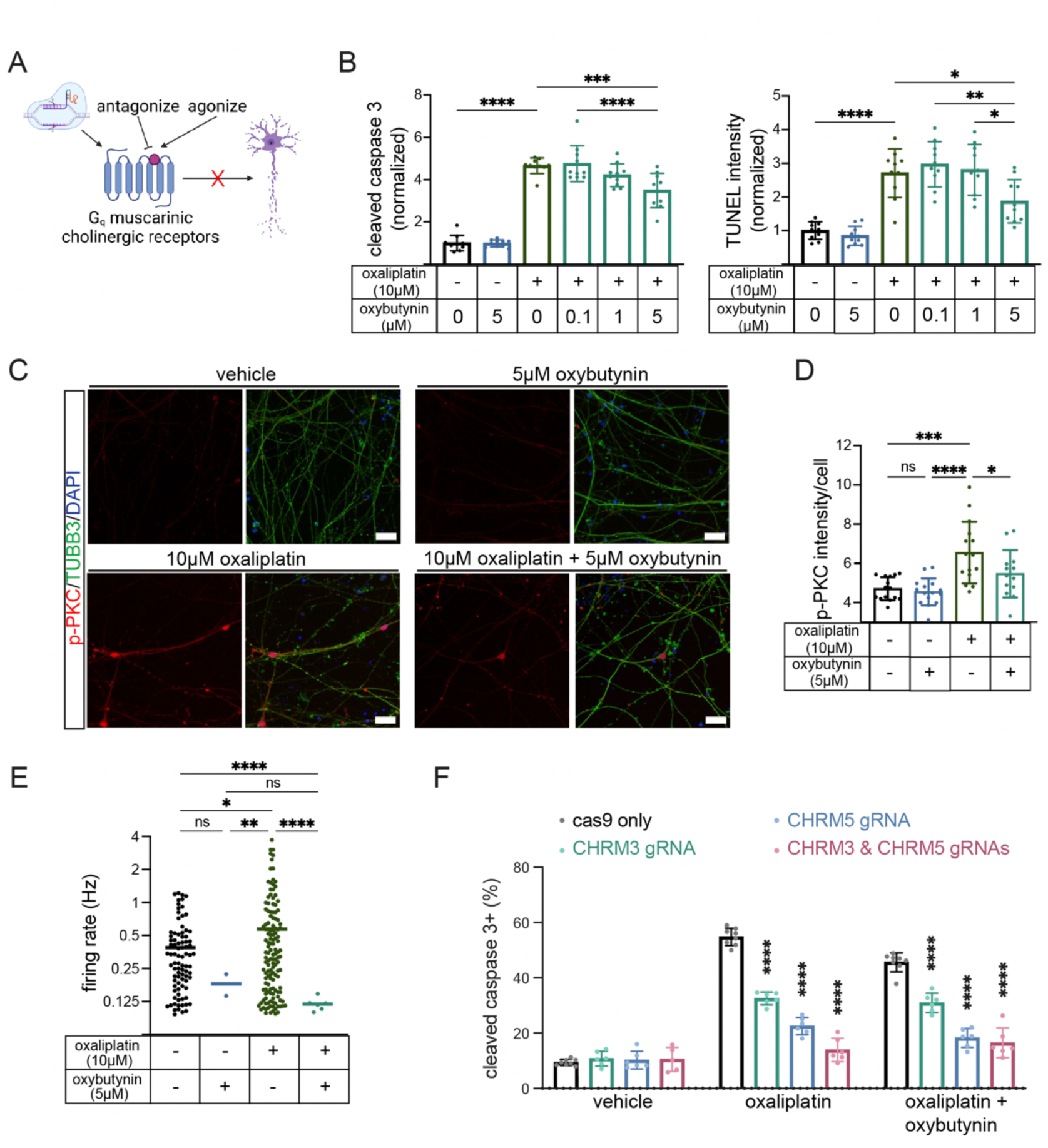
Reducing muscarinic cholinergic receptor signaling prevents platin-induced excitotoxicity in enteric neurons. **A)** Schematic illustration of our strategy to use tool compounds and CRISPR-cas9 technology to interrogate the role of muscarinic cholinergic receptor signaling in platin-induced enteric neuropathy. **B)** Quantification of image based analysis detecting the dose response of the oxybutynin on hPSC-derived enteric neuron cleaved caspase 3 (left) and TUNEL (right) levels when dosed in combination with oxaliplatin. All values are normalized to vehicle (medium containing 1% water). **C, D**) Phosphorylated protein kinase C (p-PKC) staining and quantification in oxybutynin and oxaliplatin-treated hPSC-derived enteric neurons. Vehicle condition contains water diluted to 1% in medium. **E)** Quantification of microelectrode array analysis of oxybutynin and oxaliplatin-treated hPSC-derived enteric neurons. Vehicle condition contains water diluted to 1% in medium. **F)** Quantification of flow cytometry analysis detecting cleaved caspase 3 levels during oxybutynin and oxaliplatin treatment in *CHRM3* and *CHRM5* CRISPR-cas9 RNP targeted hPSC-derived enteric neurons. Vehicle condition contains water diluted to 1% in medium. *p value < 0.05, ** p value < 0.01, *** p value < 0.001, **** p value < 0.0001, ns: not significant.

To further evaluate the ability of oxybutynin to protect against platin-induced enteric neuropathy, we measured its effect on oxaliplatin-induced apoptosis. Treatment with oxaliplatin significantly increased the percentage of cells positive for cleaved caspase 3, and oxybutynin was able to significantly reduce this effect in a dose-dependent manner (Figure 3B). We also measured the level of DNA fragmentation, a hallmark of late stage apoptosis, by terminal deoxynucleotidyl transferase dUTP nick end labeling (TUNEL) assay, and detected a significant increase upon oxaliplatin treatment. Again, oxybutynin co-treatment was able to significantly reduce this effect in a dose-dependent manner (Figure 3B). Thus, oxybutynin is highly effective at preventing oxaliplatin-induced apoptosis in enteric neurons.

To determine if oxybutynin can prevent overactivity of the inositol phospholipid signaling pathway, we detected p-PKC levels as a hallmark of pathway activity. As expected, oxaliplatin significantly increased levels of p-PKC per cell, validating increased PKC activity as a hallmark of platin-induced excitotoxicity (Figure 3C-D, S3A). When the cells were co-treated with oxybutynin and oxaliplatin, however, p-PKC levels were significantly reduced (Figure 3C-D, S3A). This suggests that antagonizing cholinergic signaling in enteric neurons is sufficient to reduce G_q_-mediated inositol phospholipid signaling pathway activity.

To determine if oxybutynin can prevent platin-induced excitotoxicity, we performed a microelectrode array experiment measuring the mean firing rate of hPSC-derived enteric neurons treated with oxaliplatin with and without oxybutynin. After three hours of incubation, oxaliplatin significantly increased the electrical firing of enteric neurons as compared to vehicle-treated enteric neurons (Figure 3E). Oxybutynin co-treatment, however, completely blocked this oxaliplatin-induced hyperexcitability, suggesting platin-induced excitotoxicity can be prevented by oxybutynin.

To determine if oxybutynin’s efficacy is mediated through muscarinic cholinergic antagonism and validate the role of muscarinic cholinergic signaling in platin-induced enteric neuropathy, we employed two structurally distinct compounds: pilocarpine, a muscarinic cholinergic receptor agonist, and solifenacin, a muscarinic cholinergic receptor antagonist. Dosing these compounds in combination with oxaliplatin produced opposite effects on oxaliplatin-induced apoptosis. Pilocarpine exacerbated toxicity, increasing the percentage of enteric neurons positive for cleaved caspase 3 by 3.2%, and solifenacin dampened toxicity, reducing the percentage of neurons positive for cleaved caspase 3 by 3.5% (Figure S3B). These results suggest that muscarinic cholinergic signaling potentiates platin-induced enteric neurotoxicity and oxybutynin’s protective effect is mediated by antagonizing this pathway.

Oxybutynin is most potent at antagonizing CHRM3 but at physiologically relevant concentrations can block all five muscarinic receptors. CHRM1, CHRM3, and CHRM5 are GPCRs of the G_q_ class, thus all could be mediating the activation of the inositol phospholipid signaling pathway involved in the platin-induced excitotoxicity mechanism. To determine which of the three receptors are expressed in enteric neurons, we measured the percentage of cells positive for CHRM1, CHRM3, and CHRM5, and found that CHRM3 and CHRM5 are the most abundant, whereas CHRM1 is expressed at negligible levels in both ESC and iPSC-derived enteric neurons (Figure S3C).

To understand which of the two expressed receptors are driving oxybutynin’s rescue effect, we targeted ESCs with *CHRM3* and *CHRM5* CRISPR-cas9 ribonucleoproteins (RNPs). Gene editing was effective at reducing targeted protein levels in differentiated enteric neurons (Figure S3D). We did not observe any compensation affecting cholinergic receptor levels in the targeted cells (Figure S3E). Furthermore, the targeted lines recapitulate the representation of key enteric neuron subtype markers, indicating that gene editing did not affect differentiation quality or lineage trajectories (Figure S3F). Treatment with *CHRM3* and *CHRM5* RNPs significantly reduced the percentage of cells positive for cleaved caspase 3 in response to oxaliplatin exposure (Figure 3F). Furthermore, oxybutynin co-treatment in the *CHRM3* and *CHRM5* targeted lines did not significantly reduce apoptosis levels further, indicating that both proteins are downstream of oxybutynin (Figure 3F).

Altogether, our results indicate that platin-induced hyperexcitability and enteric neurotoxicity is mediated though CHRM3 and CHRM5 signaling, and oxybutynin can counteract these toxic effects by antagonizing these receptors.

### Enteric nitrergic neurons are selectively vulnerable to platin-induced excitotoxicity

Enteric neurons are heterogeneous, composed of a diverse array of molecularly, structurally, and functionally distinct neuronal subtypes^7,8^. Furthermore, many enteric neuropathies, also known as disorders of gut-brain interaction, are caused by the loss or dysfunction of a specific enteric neuron subtype^38,39^. Therefore, we sought to determine how our platin-induced excitotoxicity pathway may be represented in different enteric neurons and how muscarinic cholinergic receptor antagonism may help protect them (Figure 4A). Thus, we performed a single nuclei RNA sequencing (snRNA-seq) experiment on our hPSC-derived enteric neurons treated with oxaliplatin, with and without oxybutynin to capture the cell-type specific transcriptional effects of these drugs. Dimensionality reduction and unbiased clustering revealed 10 transcriptionally distinct cell type clusters present in our cultures that do not cluster based on treatment condition (Figure S4A). Based on the expression of respective lineage markers, clusters 1, 2, 3, 5, 7, and 8 were annotated as progenitor populations, cluster 4 as enteric neurons, cluster 6 as enteric glia, cluster 8 as enteric neural crest cells, and cluster 9 as smooth muscle cells, as defined based on the expression of their respective lineage markers (Figure S4B).

**Figure 4:**
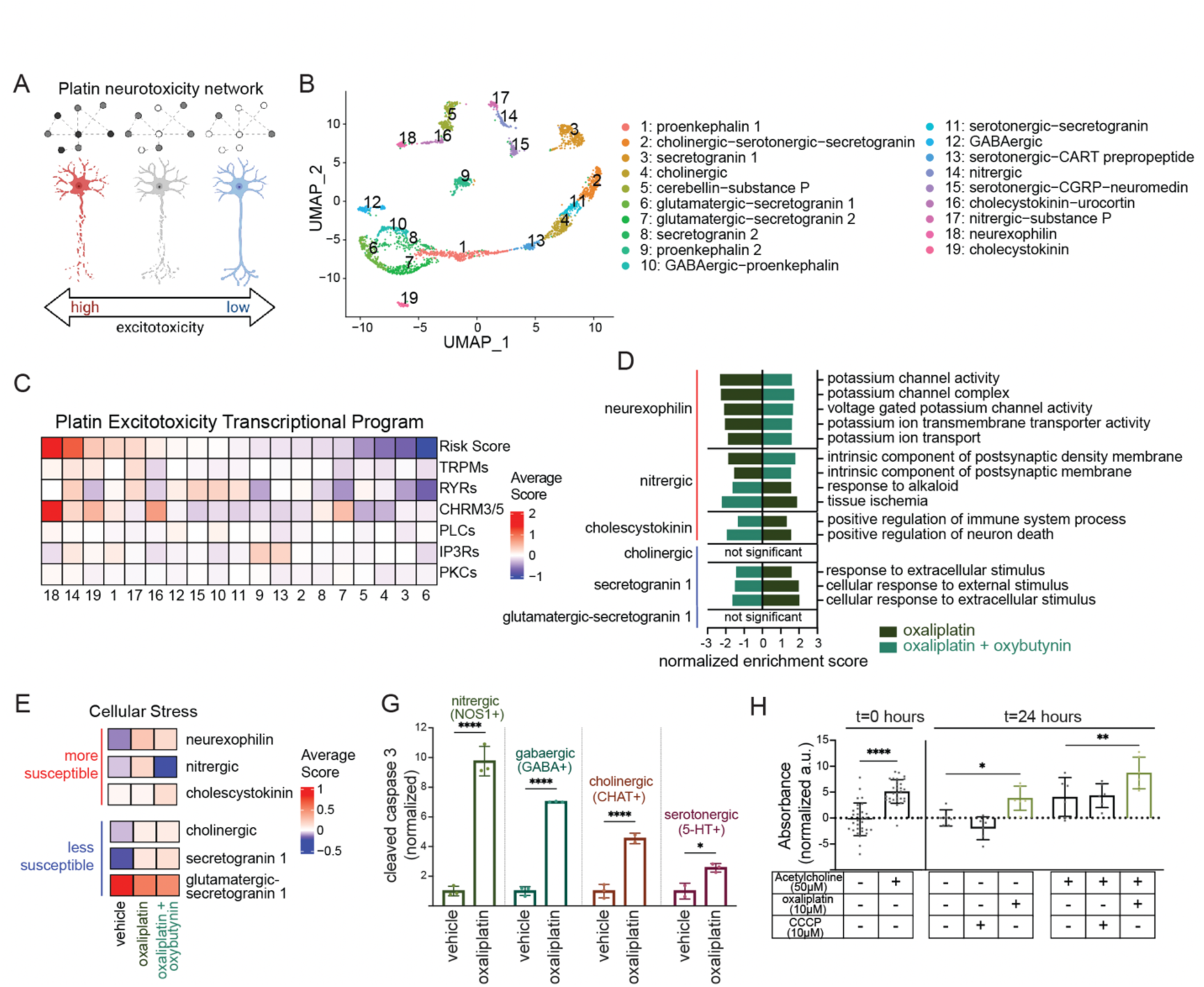
Single nuclei transcriptomic profiling identifies subtypes susceptible to platin-induced excitotoxicity. **A)** Schematic illustration of our hypothesis that enteric neuron subtypes are differentially susceptible to platin toxicity based on their expression of platin-induced excitotoxicity pathway genes. **B)** sn-RNAseq UMAP of enteric neuron subtypes present in our stage 1 2D enteric neuron cultures. **C)** Heatmap of the average module scores of the platin excitotoxicity transcriptional program gene categories in the stage 1 2D enteric neuron subtypes. **D)** Bar plot of the Ontology neurotransmission and cellular stress related gene sets rescued by oxybutynin co-treatment in the stage 1 2D enteric neuron subtypes predicted to be most susceptible (neurexophilin, nitrergic, and cholescystokinin) and least susceptible (cholinergic, secretogranin 1, and glutamatergic-secretogranin 1) to platin-induced excitotoxicity. **E)** Heatmaps of the average module scores of the combined list of all genes in the Hallmark gene sets (combined cellular stress score) in the stage 1 2D enteric neuron subtypes predicted to be most susceptible (neurexophilin, nitrergic, and cholescystokinin) and least susceptible (cholinergic, secretogranin 1, and glutamatergic-secretogranin 1) to platin-induced excitotoxicity. **F)** Quantification of flow cytometry analysis detecting cleaved caspase 3 levels during 20 μM oxaliplatin treatment in four enteric neurotransmitter identities: nitrergic (NOS1+), GABAergic (GABA+), cholinergic (CHAT+), and serotonergic (5-HT+) stage 1 2D enteric neurons. Vehicle contains water diluted to 2% in medium. Data are represented as mean ± SD. **G)** Quantification of calorimetry-based method for detecting nitric oxide levels with and without acetylcholine stimulation in the stage 1 2D enteric neuron cell culture supernatants prior to drug treatment (t=0 hours) and after 24 hours exposure to oxaliplatin and CCCP. Data is normalized by subtracting the vehicle (medium containing 1% water) pre-treatment (t=0 hours) condition from each sample. Data are represented as mean ± SD. *p value < 0.05, ** p value < 0.01, **** p value < 0.0001, ns: not significant.

To validate that our hPSC-derived enteric neurons recapitulate the identity of human enteric neurons *in vivo*, we compared them to a snRNA-seq dataset of the primary human colon previously published by Regev and colleagues^20^. Module scoring of the transcriptional signatures of the primary cell types on our hPSC-derived cell types demonstrated that the enteric neuron populations are highly similar, with our enteric neuron cluster 4 showing the highest average expression of primary human enteric neuron genes (Figure S4C). The reverse analysis, scoring the expression of our stage 1 2D enteric neuron genes in the primary human dataset, validated these findings, resulting in the primary enteric neurons having the highest average expression of our hPSC-derived enteric neuron genes (Figure S4C). Thus, our *in vitro* hPSC-derived enteric neurons share the same transcriptional identity as their *in vivo* counterparts.

To identify the neuronal subtypes represented in our hPSC-derived enteric neurons, we sub-clustered the neuronal population. Sub-clustering revealed 19 transcriptionally distinct enteric neuron clusters which were annotated by neurochemical identity based on their expression of neurotransmitter processing genes and neuropeptides (Figure 4B and S4D). Cells from the drug treatment conditions appear to cluster by neuronal subtype identity, as all 19 subtypes are represented in each of the three treatment conditions (Figure S4E).

Next, we set out to determine if our hPSC-derived enteric neurons (henceforth referred to as “stage 1 2D enteric neurons” based on their differentiation stage and format) contain the same neuronal diversity as other published enteric neuron datasets. We compared the neurochemical composition of our dataset to hPSC-derived stage 1 and stage 2 ganglioid (3D) enteric neurons published by our group^21^, primary fetal human enteric neurons published by Teichmann and colleagues^40^, as well as primary adult human colonic enteric neurons published by Regev and colleagues^20^. For neurotransmitters, we utilized a two-step neurochemical identity-defining approach, that our group has reported previously, to identify all the cells that belong to six neurotransmitter identities: serotonergic, GABAergic, catecholaminergic, glutamatergic, cholinergic, and nitrergic^21^. We found that all six neurochemical identities are represented in our cultures, recapitulating the neurotransmitter diversity of other datasets (Figure S4F). Furthermore, enteric neurons that produce multiple neurotransmitters have recently been characterized in some detail^21^. To determine if our stage 1 2D enteric neurons recapitulate the same neurotransmitter complexity as other enteric neuron tissues, we quantified the number of cells predicted to synthesize only one, two, three, or more neurotransmitters across all five datasets. Here neurotransmitter complexity was also reproduced, with our stage 1 2D enteric neurons containing cells with all four levels of complexity (Figure S4G). Next, we characterized the representation and complexity of neuropeptides and identified 15 identities based on the expression of neuropeptide genes including calcitonin 2, CART prepropeptide, cerebellin 2, cholecystokinin, chromogranin A, endothelin 3, galanin, neuromedin U, neuropeptide Y, neuroexophilin 2, proenkephalin, secretogranin 3, somatostatin, substance P, and urocortin. All five datasets have representation from all 15 neuropeptide identities with the exception of the stage 2 ganglioid enteric neurons, which only have 14, lacking CAPT prepropeptide expressing cells likely due to a sampling issue (Figure S4H). Detecting neuropeptide complexity in each dataset based on the number of cells expressing one, two, three, or more neuropeptides again revealed that different levels of neuropeptide complexity are represented across all five datasets (Figure S4I). Thus, our stage 1 2D enteric neurons recapitulate the neurochemical representation and complexity of other enteric neuron datasets, including hPSC-derived enteric ganglioids and primary human enteric neurons.

To predict which enteric neuron subtypes are more susceptible to the platin-induced excitotoxicity mechanism, we performed module scoring on our vehicle-treated enteric neuron subtypes for their expression of genes contributing to various steps in the platin-induced excitotoxicity pathway, which we term the platin excitotoxicity transcriptional program. Transcripts in this program include transient receptor potential channels, ryanodine receptors, CHRM3 and CHRM5, phospholipase C enzymes, IP3 receptors, and protein kinase C enzymes (Supplementary Table 7 and Figure 4C). We module scored the neurons using a combined transcript list to generate an overall risk score for each of the enteric neuron subtypes (Figure 4C). The neurexophilin, nitrergic, and cholescystokinin populations were predicted to be the most susceptible enteric neuron subtypes. The neurexophilin population’s high-risk score is primarily driven by high relative expression of *CHRM3* and *CHRM5*, whereas nitrergic neurons have high relative expression of all six gene categories (Figure 4C). Although neurexophilin and cholescystokinin enteric neurons have been identified in the ENS and appear to have a transcriptional profile associated with intrinsic primary afferent neurons, their functional role in the ENS and association with enteric neuropathies has been understudied^41–43^. Nitrergic neurons, on the other hand, are better characterized, releasing nitric oxide to relax smooth muscle cells in the GI tract, thereby promoting GI motility^9^. Enteric neuropathies associated with nitrergic neurons include achalasia, hypertrophic pyloric stenosis, and gastroparesis^9^. Furthermore, in a mouse model of oxaliplatin-induced enteric neuropathy, nitrergic neuron-mediated electrophysiological activities and motility functions are disrupted, which aligns with our data, predicting them to be more susceptible to oxaliplatin-induced excitotoxicity^14^. The cholinergic, secretogranin 1, and glutamatergic-secretogranin 1 populations, on the other hand, were predicted to be the least susceptible enteric neuron subtypes with low expression of virtually every gene category involved in the platin-induced excitotoxicity pathway (Figure 4C).

Next, we assessed if our predictions are reflected in the transcriptional changes induced by the drug treatments in each of the subtypes. To do this, we performed pre-ranked GSEA on the genes differentially expressed between vehicle and oxaliplatin-treated neurons and oxaliplatin and oxaliplatin-oxybutynin co-treated neurons for the neurexophilin, nitrergic, cholecystokinin, cholinergic, secretogranin 1, and glutamatergic-secretogranin 1 populations. We filtered the significant gene sets using a p-value cutoff of < 0.05, then identified the gene sets with opposite normalized enrichment scores for each of the comparisons per population, indicating gene sets rescued by oxybutynin co-treatment. Focusing on the gene sets relating to neurotransmission and cellular stress pathways, we identified five that were rescued for neurexophilin neurons, four for nitrergic neurons, two for cholecystokinin neurons, three for secretogranin 1 neurons, and none for cholinergic neurons or glutamatergic-secretogranin 1 neurons (Figure 4D). Many of the reversed genesets were related to neuronal activity, synaptic function, and cellular stress (Figure 4D).

To validate these findings, we module scored the neurexophilin, nitrergic, cholecystokinin, cholinergic, secretogranin 1, and glutamatergic-secretogranin 1 populations for their expression of the rescued neurotransmission and cellular stress related gene sets. Comparing the average relative expression of each module across each population’s treatment conditions, we observed that the neurexophilin, nitrergic, and cholecystokinin subtypes exhibit expression patterns that are partially or fully reversed (Figure S5A). This pattern is not recapitulated in the cholinergic, secretogranin 1, and glutamatergic-secretogranin 1 subtypes where only one or two of the gene sets are modulated by oxaliplatin and rescued by oxybutynin (Figure S5A). All together, these analyses indicate that neurotransmission and cellular stress related pathways in cholinergic, secretogranin 1, and glutamatergic-secretogranin 1 enteric neurons are less affected by oxaliplatin chemotherapy than they are in the neurexophilin, nitrergic, and cholecystokinin enteric neuron subtypes. Furthermore, oxybutynin can largely reverse these oxaliplatin-induced neurotransmission and cellular stress related transcriptional changes, particularly in the subtypes predicted to be more susceptible to platin-induced excitotoxicity.

To more robustly characterize the cellular stress related transcriptional changes induced by the drug treatments in the subtypes predicted to be more susceptible and less susceptible to platin-induced excitotoxicity, we module scored the neurexophilin, nitrergic, cholecystokinin, cholinergic, secretogranin 1, and glutamatergic-secretogranin 1 populations for their expression of all the Hallmark cellular stress related gene sets. These gene lists were combined, and all the neurons were module scored for the combined gene list to generate an overall cellular stress score for each of the populations. The average relative expression of the cellular stress transcriptional signature was increased by oxaliplatin and decreased by oxybutynin co-treatment for both the neurexophilin and nitrergic subtypes (Figure 4E). This pattern was not reflected in the cholinergic, secretogranin 1, and glutamatergic-secretogranin 1 subtypes, where the average relative expression of the cellular stress module increased slightly for the cholinergic and secretogranin 1 neurons and decreased slightly for the glutamatergic-secretogranin 1 neurons (Figure 4E). This cellular stress expression pattern of rescue for the two subtypes predicted to be the most susceptible to our platin-induced excitotoxicity mechanism is largely explained by eight of the cellular stress related Hallmark gene sets, including transcriptional changes related to DNA damage, oxidative stress, apoptosis, and inflammation (Figure S5B). These pathways overlap with the cellular stress hallmarks we identified through bulk RNA sequencing and follow-up experimental validations (Figure 1, Figure S1D), suggesting that the transcriptional and phenotypic changes we observed may be largely explained by the subtypes we predicted to be more susceptible to platin-induced excitotoxicity.

We were intrigued by the prediction that nitrergic neurons (clusters 14 and 17) are more susceptible to platin-induced excitotoxicity and cholinergic neurons (clusters 2 and 4) are less susceptible (Figure 4C), since proper GI motility is maintained through a balance of cholinergic (excitatory) and nitrergic (inhibitory) motor neuron signaling^7,8^. Cholinergic neurons release acetylcholine to promote the contraction of smooth muscle cells that drive GI motility and nitrergic neurons are responsible for inducing muscle relaxation through the release of nitric oxide^7,8^. So, a selective loss of nitrergic neurons following platin chemotherapy treatment could lead to excessive contraction of the smooth muscle tissue contributing to the dysfunctional GI symptoms experienced by patients (Figure S1B, Supplementary Table 3). To continue interrogating this hypothesis, we further characterized the vehicle-treated enteric neurons, detecting the expression of the plain excitotoxicity transcriptional program genes in nitrergic and cholinergic neurons relative to all the other enteric neurons in our dataset. We defined the nitrergic neuron clusters based on the high percentage of cells expressing the nitric oxide rate limiting enzyme *NOS1* (Figure S4D). However, some *NOS1* expressing cells can be found in other clusters (Figure S4D). Similarly, cholinergic neurons, expressing the acetylcholine rate limiting enzyme *CHAT*, exist outside of the cholinergic neuron clusters (Figure S5C). To take a more holistic look at these two these populations, we identified all nitrergic neurons and cholinergic neurons using our two-step neurochemical identity-defining approach^21^, thereby binning all enteric neurons in our dataset into nitrergic, cholinergic, or other categories (Figure S5C). We then module scored the nitrergic, cholinergic, and other neurons based on their expression of the genes in the platin excitotoxicity transcriptional program (Supplementary Table 7). Nitrergic neurons have a higher relative expression of the platin-induced excitotoxicity related genes relative to other enteric neurons, whereas cholinergic neurons have a lower relative expression of these same genes relative to other enteric neurons (Figure S5D). These data further support our prediction that nitrergic neurons are more susceptible and cholinergic neurons are less susceptible to platin-induced excitotoxicity.

To experimentally validate if nitrergic neurons are indeed more susceptible to platins than other enteric neuron subtypes, we detected levels of oxaliplatin-induced cleaved caspase 3 across four enteric neurotransmitter identities: CHAT+ cholinergic neurons, GABA+ gabaergic neurons, NOS1+ nitrergic neurons, and 5-HT+ serotonergic neurons. Apoptosis was induced to different degrees across these populations, with the nitrergic population showing the strongest response, with apoptosis levels increasing nearly 10-fold relative to vehicle-treated nitrergic neurons (Figure 4F). Apoptotic cholinergic neurons, on the other hand, only increased 4.5-fold relative to vehicle levels (Figure 4F). These results confirm that nitrergic neurons are more susceptible to oxaliplatin relative to other enteric neuron populations.

To validate that the increased susceptibility of nitrergic neurons to oxaliplatin is a consequence of oxaliplatin-induced hyperexcitability, we measured the levels of nitric oxide released by the nitrergic neurons as a result of their electrical activity. Stimulation with acetylcholine leads to an increase in nitric oxide release by the nitrergic neurons (Figure 4G). This result functionally validates that nitrergic neurons are responsive to acetylcholine, as predicted based on their expression of *CHRM3* and *CHRM5* (Figure 4C and Figure S5D). We then exposed our enteric neuron cultures to oxaliplatin for 24 hours and measured the levels of nitric oxide released into the medium in the presence and absence of acetylcholine stimulation. In both conditions, we observed that oxaliplatin-treated nitrergic neurons released significantly more nitric oxide than their vehicle-treated counterparts. This effect was not observed when the cells were treated with the mitochondrial toxin CCCP, indicating that the effect is specific to oxaliplatin toxicity and not due to a general susceptibility of nitrergic neurons to cellular stress (Figure 4G).

Altogether, our results indicate that enteric neuron subtypes are differentially susceptible to platin chemotherapies. We demonstrated that nitrergic neurons show a stronger apoptotic response relative to other neurons. Furthermore, this increased level of cell death is preceded by hyperexcitability of nitrergic neurons and increased release of nitric oxide. These data suggest that preserving the susceptible neuronal subtypes by inhibiting cholinergic signaling, might be a promising therapeutic approach to prevent the clinical symptoms of platin-induced GI neurotoxicity.

### Oxybutynin prevents oxaliplatin-induced enteric neuropathy in mice

Our results suggest that cholinergic signaling, mediated through CHRM3 and CHRM5, increases the sensitivity of enteric neurons to platins via an excitotoxicity mechanism, which occurs in a subtype-specific manner, based on the expression of the platin excitotoxicity transcriptional program. To determine if hallmarks of platin-induced excitotoxicity are represented in platin chemotherapy patient tissue, we acquired archived histologically normal gastric tissue from prior gastrectomies for gastric cancer, that either did not receive chemotherapy prior to tumor resection or did undergo platin chemotherapy prior to tumor resection (Figure 5A). We first evaluated if platin chemotherapy treatment increases activity of the inositol phospholipid signaling pathway, a key component of the platin excitotoxicity mechanism, by detecting p-PKC levels by immunostaining. Similar to what we observed *in vitro*, patients that had received platin chemotherapy prior to their surgery had elevated p-PKC levels in their enteric neurons relative to the patients that had not undergone chemotherapy (Figure 3C-D, Figure S3A, Figure 5B-C). Thus, these data support our findings that platins increase inositol phospholipid signaling and PKC activity in human enteric neurons. To evaluate if platin chemotherapy treatment alters the representation of the enteric neuron subtypes we identified as more or less susceptible to platin-induced excitotoxicity, we detected nitrergic neurons and cholinergic neurons based on the colocalization of the pan-neuronal marker HuC/D with NOS1 and CHAT respectively by immunostaining (Figure 5D). We quantified the proportion of NOS1+ CHAT-, CHAT+ NOS1-, and CHAT+ NOS1+ enteric neurons in each patient group (Figure 5E). The proportion of NOS1+ CHAT-enteric neurons was significantly reduced, while the proportion of CHAT+ NOS1-enteric neurons was significantly increased in patients that received platin chemotherapy prior to surgery relative to those that did not undergo chemotherapy (Figure 5E). These data support our *in vitro* observations that nitrergic enteric neurons are selectively vulnerable to platin chemotherapies as compared to the more resistant cholinergic subtype.

**Figure 5:**
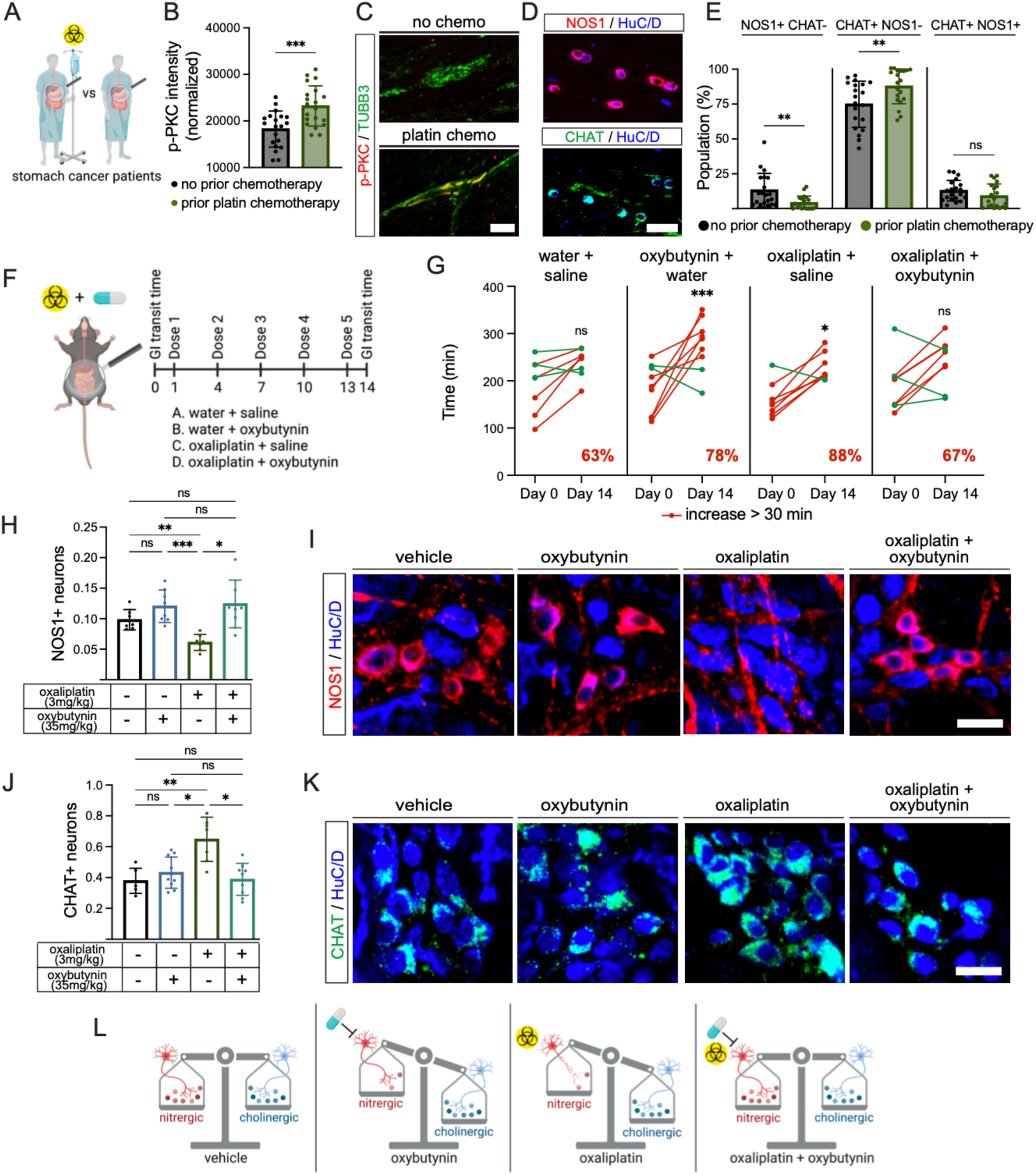
Selective loss of nitrergic enteric neurons during platin treatment can be prevented by oxybutynin. **A)** Schematic illustration of our strategy to stratify gastric cancer patients to compare by immunostaining the enteric neurons of patients that received platin chemotherapy to patients that received no chemotherapy prior to their tumor resection surgeries. All staining was done in non-cancerous healthy tissue margins from the surgical samples. **B, C)** Immunofluorescence staining and quantification of primary human stomach sections for protein kinase C activity marker phosphorylated protein kinase C (p-PKC) and neuronal marker TUBB3 comparing patients that received platin chemotherapy to patients that received no chemotherapy prior to their tumor resection surgeries. Data are represented as mean ± SD. Scale bar: 50 μm. **D)** Immunofluorescence staining of primary human stomach sections for nitrergic neuron marker NOS1 and cholinergic neuron marker CHAT. Scale bar: 25 μm. **E)** Bar plot showing the proportion of NOS1+ CHAT-nitrergic neurons, CHAT+ NOS1-cholinergic neurons, and CHAT+ NOS1+ dual identity enteric neurons in primary human stomach sections of patients that received platin chemotherapy to patients that received no chemotherapy prior to their tumor resection surgeries. Data are represented as mean ± SD. **F)** Schematic illustrating the design of our mouse study where mice were dosed with oxaliplatin (3 mg/kg per dose), oxybuytnin (35 mg/kg per dose), oxaliplatin’s vehicle (3 mg/kg water per dose), and oxybutynin’s vehicle (35 mg/kg 5% DMSO and 2% tween80 in saline). GI transit time was measured before and after the dosing regimen. Enteric neurons were then compared between the different treatment conditions by immunostaining. **G)** Paired analysis of GI transit times for each animal by treatment condition comparing GI transit times of the animals before and after the dosing regimen. Red lines represent mice that experienced a > 30 minute increase in their GI transit. Green lines represent mice that experienced < 30 minute increase in their GI transit. Percentages reflect the percentage of mice per condition that experienced a > 30 minute increase in their GI transit. **H, I)** Nitrergic neuron marker NOS1 and neuronal nuclei marker (HuC/D) staining and quantification in vehicle (3 mg/kg water + 35 mg/kg 5% DMSO and 2% tween80 in saline), oxybuytnin (35 mg/kg oxybutynin + 3 mg/kg water), oxaliplatin (3 mg/kg oxaliplatin + 35 mg/kg 5% DMSO and 2% tween80 in saline), and oxaliplatin + oxybutynin (3 mg/kg oxaliplatin + 35 mg/kg oxybutynin)-treated mice. Data are represented as mean ± SD. Scale bar: 25 μm. **J, K)** Cholinergic neuron marker CHAT and neuronal nuclei marker (HuC/D) staining and quantification in vehicle (3 mg/kg water + 35 mg/kg 5% DMSO and 2% tween80 in saline), oxybuytnin (35 mg/kg oxybutynin + 3 mg/kg water), oxaliplatin (3 mg/kg oxaliplatin + 35 mg/kg 5% DMSO and 2% tween80 in saline), and oxaliplatin + oxybutynin (3 mg/kg oxaliplatin + 35 mg/kg oxybutynin)-treated mice. Data are represented as mean ± SD. Scale bar: 25 μm. **L)** Schematic illustrating conclusions from the mouse study. *p value < 0.05, ** p value < 0.01, *** p value < 0.001, ns: not significant.

To determine if muscarinic cholinergic receptor antagonism, via oxybutynin, can prevent the cellular and physiological hallmarks of platin-induced GI neurotoxicity *in vivo*, we designed a mouse experiment including four treatment conditions: vehicle-treated mice injected with water and saline, oxybutynin-treated mice injected with water and oxybutynin, oxaliplatin-treated mice injected with oxaliplatin and saline, and oxaliplatin and oxybutynin co-treated mice injected with both oxaliplatin and oxybutynin (Figure 5H). All mice were injected following an established dosing regimen previously shown to induce hallmarks of oxaliplatin-induced enteric neuropathy^13,14^. The total GI transit time of each mouse was measured on days zero and 14 to assess if the treatments significantly change the time it takes for food to pass through the GI tract (Figure 5F). Consistent with previous reports^13,14^, oxaliplatin treatment led to significantly longer GI transit times, with 88% of the mice showing an increase of more than 30 minutes (Figure 5G). Oxybutynin treatment alone also significantly increased GI transit time, with 78% of the mice showing a more than 30 minute increase (Figure 5G). This result likely reflects oxybutynin’ antagonizing effect on muscarinic receptors in the ENS. Mice co-treated with oxaliplatin and oxybutynin, however, did not exhibit significant increases in GI transit time (Figure 5G). These results highlight the therapeutic potential of oxybutynin, to prevent physiological hallmarks of platin-induced GI neuropathy.

Our data suggests that antagonizing muscarinic cholinergic receptors during platin treatment can prevent clinical features of platin-induced GI neurotoxicity. We next sought to determine if the functional outcome of oxaliplatin-induced constipation in our study reflects a change in representation of neuronal subtypes in the ENS and evaluate whether oxybutynin can rescue these cellular features. We quantified the number of nitrergic and cholinergic neurons in the distal colon of the treated mice using immunostaining of the pan neuronal marker HuC/D with NOS1 and CHAT, respectively. The proportion of nitrergic neurons in the mice treated with oxaliplatin was significantly reduced, nearly in half, relative to vehicle levels (Figure 5H-I). Oxybutynin co-treatment was able to prevent the oxaliplatin-induced reduction of the nitrergic neuron population, suggesting that oxybutynin can protect this susceptible subtype from platin toxicity *in vivo* (Figure 5H-I). Conversely, the proportion of cholinergic neurons in mice treated with oxaliplatin was significantly increased relative to vehicle levels, suggesting this population is resistant to platins and, therefore, becomes overrepresented in the tissue (Figure 5J-K). Oxybutynin co-treatment was able to prevent this overrepresentation, likely through its preservation of more vulnerable enteric neuron subtypes, such as the nitrergic neurons (Figure 5J-K). Mice treated only with oxybutynin did not exhibit significant changes in their neuronal proportions relative to vehicle-treated mice, suggesting that the prolonged GI transit time exhibited by these animals reflects a reduction in nitrergic neuron activity, via muscarinic cholinergic receptor antagonism, rather than a loss of the neuronal population due to toxicity (Figure 5H-K).

Altogether, the results suggests that platin treatment disrupts the balance between nitrergic and cholinergic neuron levels and activity, required for proper GI function under normal conditions (Figure 5L). Selective vulnerability of nitrergic neurons to platins, creates an imbalance in the ENS characterized by an overrepresentation of excitatory cholinergic neurons and an underrepresentation of inhibitory nitrergic neurons, thereby causing GI dysmotility (Figure 5L). When treated alone, oxybutynin antagonizes muscarinic cholinergic receptors on inhibitory nitrergic neurons, dampens their activity and leads to slow GI transit (Figure 5L). However, in combination with oxaliplatin treatment, oxybutynin prevents platin-induced excitotoxicity, thereby preserving the vulnerable nitrergic neurons and preventing platin-induced GI dysmotility (Figure 5L). In summary, these data support muscarinic cholinergic receptor antagonism as a viable therapeutic strategy to prevent platin-induced GI neuropathy.

## Discussion

Enteric neurons innervate and control the GI tract^7,8^. Medications such as chemotherapy agents, can damage enteric neurons, causing a variety of GI functional defects, like severe constipation^13–18^. Platin chemotherapies contribute heavily to the burden of chemotherapy-induced GI neurotoxicity, causing symptoms that can be dose-limiting and irreversible, significantly impacting quality of life^12^. However, despite being on the market for decades, the cell type-specific mechanisms underlying platin-induced GI neurotoxicity have remained largely unknown.

Here, we leveraged hPSC-derived enteric neurons as a novel model system to study platin-induced enteric neuropathy. Our comprehensive phenotypic and transcriptomics profiling revealed that the enteric nerves are more sensitive to platins relative to the other peripheral nerve populations. For example, enteric neurons showed a higher cytotoxic response and neuroinflammatory signature. We demonstrated that platins induce DNA damage, oxidative stress, neurite degeneration, and apoptosis in enteric neurons. Given that all experiments were performed in *in vitro* cultures of enteric neurons, without confounding interactions with other tissues in the GI microenvironment, these data suggest that platin-induced enteric neuropathy occurs as a result of the direct effect of platins on enteric neurons. Thus, identifying and blocking the mechanism that makes enteric neurons highly sensitive to platins would be an effective therapeutic strategy to protect this vulnerable tissue.

Taking advantage of the scalability of our hPSC-derived platin-induced enteric neuropathy model system, we performed high-throughput screening to identify drugs that regulate platin-induced enteric neuron cell death. A common characteristic of the drugs from our screen was the ability to target GPCRs. Analysis of the two GPCR classes represented in the dataset revealed that antagonizing the G_q_-mediated inositol phospholipid signaling pathway can preserve enteric neuron viability during oxaliplatin treatment, suggesting platin-induced enteric neuropathy is potentiated via a G_q_ GPCR-mediated excitotoxicity mechanism. Indeed, we found that oxaliplatin causes enteric neurons to become hyperexcitable within a few hours of treatment, triggering the excitotoxicity cascade. Furthermore, treatment with our drug candidate oxybutynin, which is a potent muscarinic cholinergic receptor antagonist, blocks this G_q_ GPCR-mediated excitotoxicity through CHRM3 and CHRM5. Notably, in colorectal cancer cell lines, dosing oxybutynin in combination with platins does not affect the efficacy of platins in killing cancer cells, making oxybutynin an ideal candidate for combination therapy in the clinic.

The ability of the enteric neurons to control diverse GI function relies on the diversity of enteric neuron subtypes represented in the ENS across different regions of the gut^38,44,45^. Thus, to understand how platins affect different enteric neuron subtypes, we performed single nuclei RNA sequencing of our hPSC-derived enteric neurons treated with vehicle, oxaliplatin, and oxaliplatin-oxybutynin co-treatment. By transcriptomic profiling and further phenotypic validation, we demonstrated that nitrergic neurons are selectively vulnerable to platins. Furthermore, oxybutynin co-treatment was sufficient to reverse toxicity related transcriptional signatures specifically in the subtypes predicted to be most vulnerable. This analysis highlights the importance of using relevant cellular models to study disease mechanisms or drug toxicity. In line with the observation that antagonizing CHRM3 and CHRM5 can prevent toxicity in enteric neurons, the representation of these receptors as well as other effectors within the platin excitotoxicity transcriptional program serves as a platin sensitivity dial that either turns up or down platin-induced toxicity in specific enteric neuron subtypes. Thus, identifying the protein network responsible for toxicity within a specific cellular context can help uncover the mechanisms underlying the cell type specificity of the disease.

Platin-induced enteric neuropathy has been detected in nearly every region of the GI tract, indicating that toxicity is not region specific and that clinical manifestations of platin-induced enteric neuropathy could vary widely, from chemotherapy-induced achalasia, gastroparesis, or constipation^13–18^. Indeed, we observed that patients that received platin chemotherapy treatment prior to their tumor resection surgery showed altered representation of enteric neuron subtypes in their stomach. The nitrergic neuron population, which we identified as selectively vulnerable to platins, was significantly reduced, whereas the cholinergic neuron population, which we identified as more resistant to platins, was significantly elevated. This overrepresentation of cholinergic neurons is likely a consequence of nitrergic neurons getting selectively eliminated by platins. This observation was consistent with the results of our mouse study where oxaliplatin significantly reduced nitrergic neuron levels and significantly increased cholinergic neurons levels; oxybutynin co-treatment was able to prevent this disruption in neuronal ratio. Furthermore, we found that co-administration of oxybutynin with oxaliplatin was able to prevent oxaliplatin-induced constipation in these animals, highlighting its therapeutic potential to prevent cellular and physiological hallmarks of platin-induced GI neuropathy.

In conclusion, our hPSC-derived model of platin-induced enteric neuropathy enabled the discovery of the mechanism underlying the selective vulnerability of enteric neurons and enteric neuron subtypes to platin chemotherapies. The work implicates an excitotoxicity cascade in the pathogenesis of platin-induced enteric neuropathy and presents muscarinic cholinergic receptor antagonism as an effective therapeutic strategy to prevent excitotoxicity and enteric neuropathy both *in vitro* and *in vivo*. Lastly, this work exemplifies how hPSC technology can enable disease modeling and drug discovery for peripheral neuropathies, serving as a roadmap for uncovering the cell type-specific responses to cellular stress underlying these historically intractable disorders.

## Acknowledgements

We thank UCSF Parnassus Flow CoLab, RRID:SCR_018206, the UCSF Laboratory for Cell Analysis, and the Human Immune Monitoring Center at Stanford University-Immunoassay Team for their excellent technical support. We acknowledge the UCSF Center for Advanced Technology, UCSF Genomics CoLab, and the 10x Genomics team for high-throughput sequencing and other genomic analyses. The work was supported by grants from UCSF Program for Breakthrough Biomedical Research and Sandler Foundation, the NIH Director’s New Innovator Award (DP2NS116769) and the National Institute of Diabetes and Digestive and Kidney Diseases (R01DK121169) to F.F.

## Author Contributions

M.N.R. Designed, completed, and analyzed all *in vitro* experiments. Designed and completed computational analyses including UCSF Research Data Browser analysis, bulk RNA sequencing data analysis, drug-protein interaction analysis, protein-protein interaction network analysis, and single nuclei RNA sequencing data analysis. Designed and led the mouse experiment, including whole mount sample preparation, staining, and analysis. Designed and led the primary human stomach tissue experiment, including staining and analysis. Wrote the manuscript.

S.F. Performed the immunostaining of the paraffin-embedded human stomach sections and assisted with the mouse assays and tissue preparation.

R.M.S. Performed the phosphorylated protein kinase C distribution analysis and assisted with the single nuclei RNA sequencing data analysis.

H.M. Assisted with *in vitro* experiments, protein-protein interaction network analysis, preparation of paraffin-embedded human stomach sections, the mouse assays, and mouse tissue preparation.

A.K.C. Assisted with the mouse assays and tissue preparation.

J.T.R. Assisted with mouse tissue preparation.

A.M. Assisted with the mouse assays.

M.D.S. Assisted with the mouse assays.

N.E. Assisted with genotyping the CHRM3 and CHRM5 CRISPR ribonucleoprotein targeted cell lines.

A.C. Assisted with the single nuclei RNA sequencing data analysis and mouse tissue preparation.

K.G. Performed injections for the mouse experiment and assisted with mouse assays.

T.J. Performed injections for the mouse experiment.

M.G.K. Performed microelectrode array data curation.

B.S. Assisted with colorectal cancer cell line experiments.

E.M.T. Assisted with quantifying CHRM3 and CHRM5 protein levels in CHRM3 and CHRM5 CRISPR ribonucleoprotein targeted cell lines by western blot.

J.Y. Assisted with bulk RNA sequencing library preparation.

A.A. Performed bulk RNA sequencing data curation.

K.Y. Assisted with the injections for the mouse experiment.

B.C. Assisted with acquiring and establishing cancer cell lines.

B.V. Assisted with UCSF Research Data Browser analysis.

C.X. Assisted with neurite length analysis.

M.G.K. Acquired the paraffin-embedded human stomach sections.

R.I. Assisted with the cell viability assays.

D.L.K. Supervised the neurite length analysis.

T.J.N. Supervised the microelectrode array data curation.

H.G. Supervised bulk RNA sequencing sample preparation and data curation, assisted with acquiring and establishing cancer cell lines, and supervised the mouse experiment.

F.F. Designed and conceived the study, supervised all experiments, and wrote the manuscript.

---

**Supplementary Figure 1:**
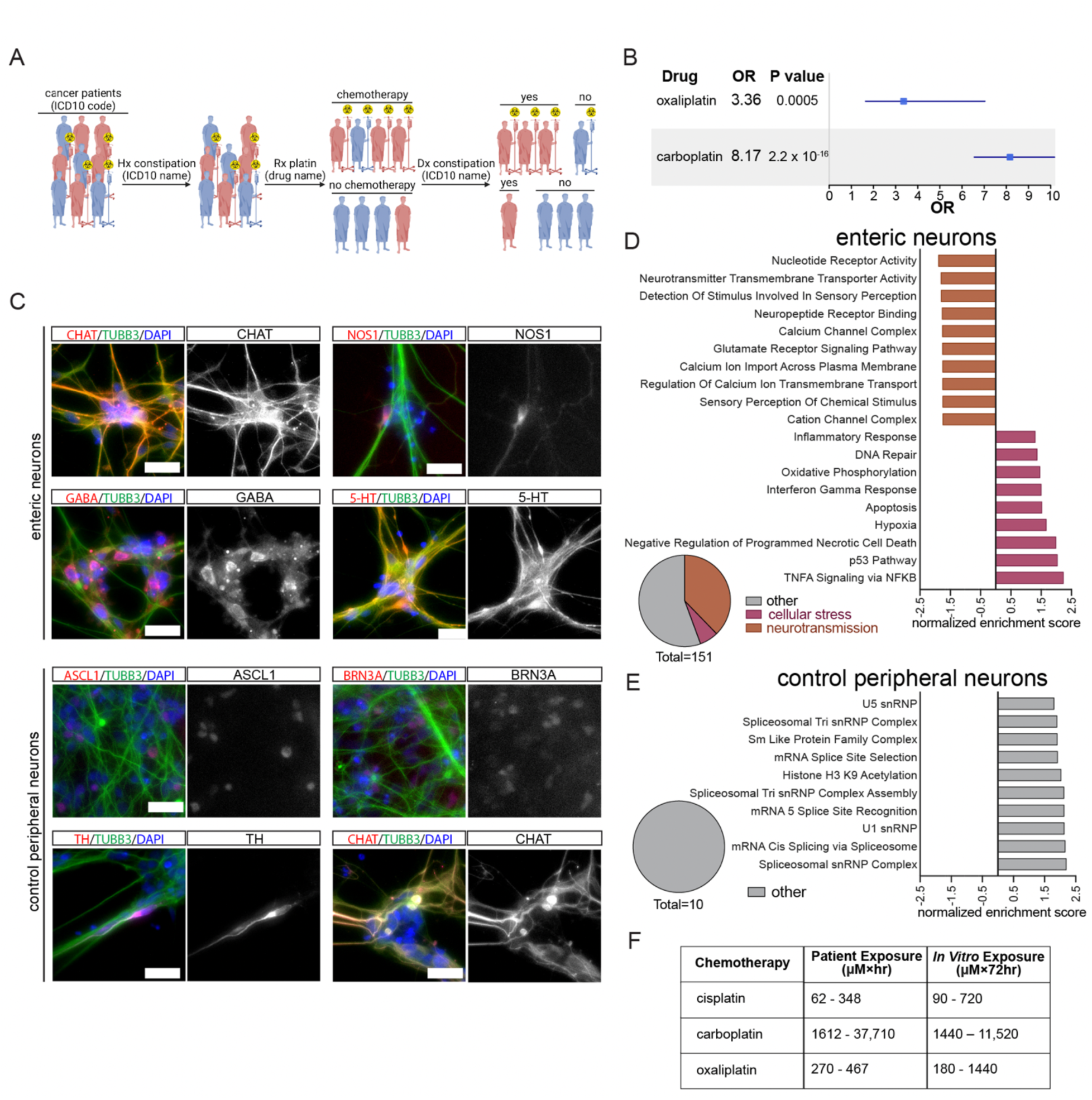
Evaluating platin-induced neurotoxicity in patient data and hPSC-derived enteric and control peripheral neurons. **A)** Schematic illustration of methods used to extract information from electronic medical records. Cancer patients were identified based on the relevant ICD10 codes. Patients with a prior history of constipation were removed based on a diagnosis of constipation (ICD10 name) prior to their cancer diagnosis. The patients were stratified depending on whether they were prescribed the relevant platin or not based on the drug name. Finally, the number of patients in each group that were diagnosed with constipation based on ICD10 name were quantified. The same analysis pipeline was used for both the oxaliplatin and carboplatin analyses. **B)** Odds ratio (OR) and p-values comparing the prevalence of constipation in patients that received platin chemotherapy to patients that did not. **C)** Representative images of enteric (CHAT, NOS1, GABA, 5-HT), sensory (BRN3A), sympathetic (ASCL1, TH), and parasympathetic (CHAT) markers in hPSC-derived enteric and control peripheral neurons. Scale bar: 50 μm. **D)** Summary of gene sets significantly enriched in the log_2_ fold-change gene expression list comparing 50 μM oxaliplatin-treated enteric neurons to vehicle (medium containing 5% water)-treated enteric neurons. The bar graph highlights the top neurotransmission and cellular stress related pathways, and the pie chart shows the proportion of significantly enriched gene sets relating to neurotransmission or cellular stress. **E)** Summary of gene sets significantly enriched in the log_2_ fold-change gene expression list comparing 50 μM oxaliplatin-treated control peripheral neurons to vehicle (medium containing 5% water)-treated control peripheral neurons. The bar graph highlights the gene sets significantly enriched in the gene list. **F)** Table including calculated platin patient exposures and hPSC-derived enteric neuron exposures. Patient exposures were calculated based on each drug’s reported clearance and a 1.79 m^2^ patient body surface area for each indication the drugs are approved for. *In vitro* exposures were calculated by multiplying the drug treatment concentrations by the duration of the experiment (72 hours).

**Supplementary Figure 2:**
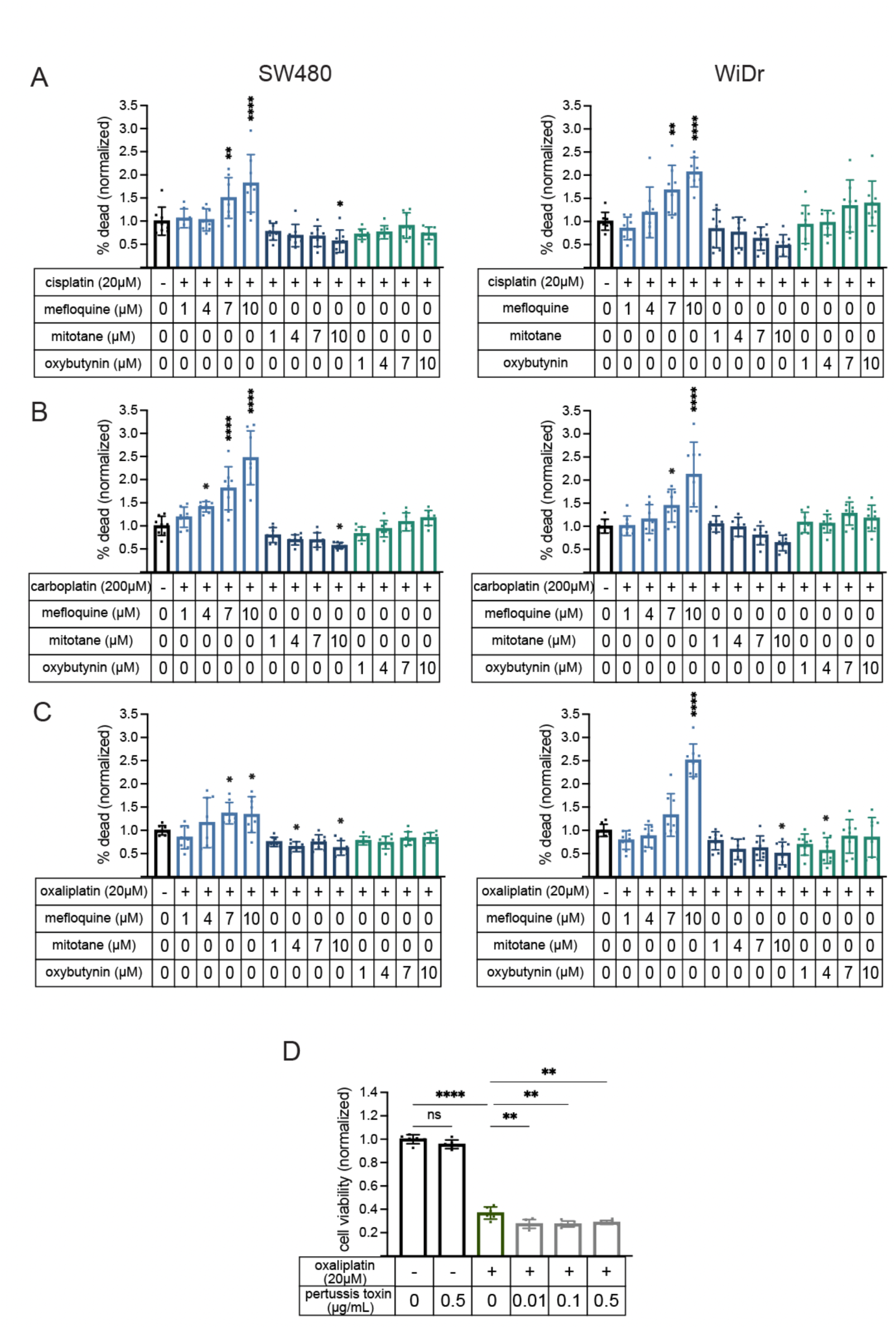
Effect of drug screen hits on platin antineoplastic efficacy and Gi protein inhibitor on platin-induced enteric neuropathy. **A-C)** Dose response of the hit compounds on SW480 (left) and WiDr (right) colorectal cancer cell viability when dosed in combination with platins. All values are normalized to the platin being tested. (A) 20 μM cisplatin, (B) 200 μM carboplatin, (C) 20 μM oxaliplatin. Data are represented as mean ± SD. **D)** Dose response of the pertussis toxin, a G_i_ inhibitor, on hPSC-derived enteric neuron viability when dosed in combination with oxaliplatin. All values are normalized to vehicle (medium containing 2% water). Data are represented as mean ± SD. *p value < 0.05, ** p value < 0.01, *** p value < 0.001, **** p value < 0.0001, ns: not significant.

**Supplementary Figure 3:**
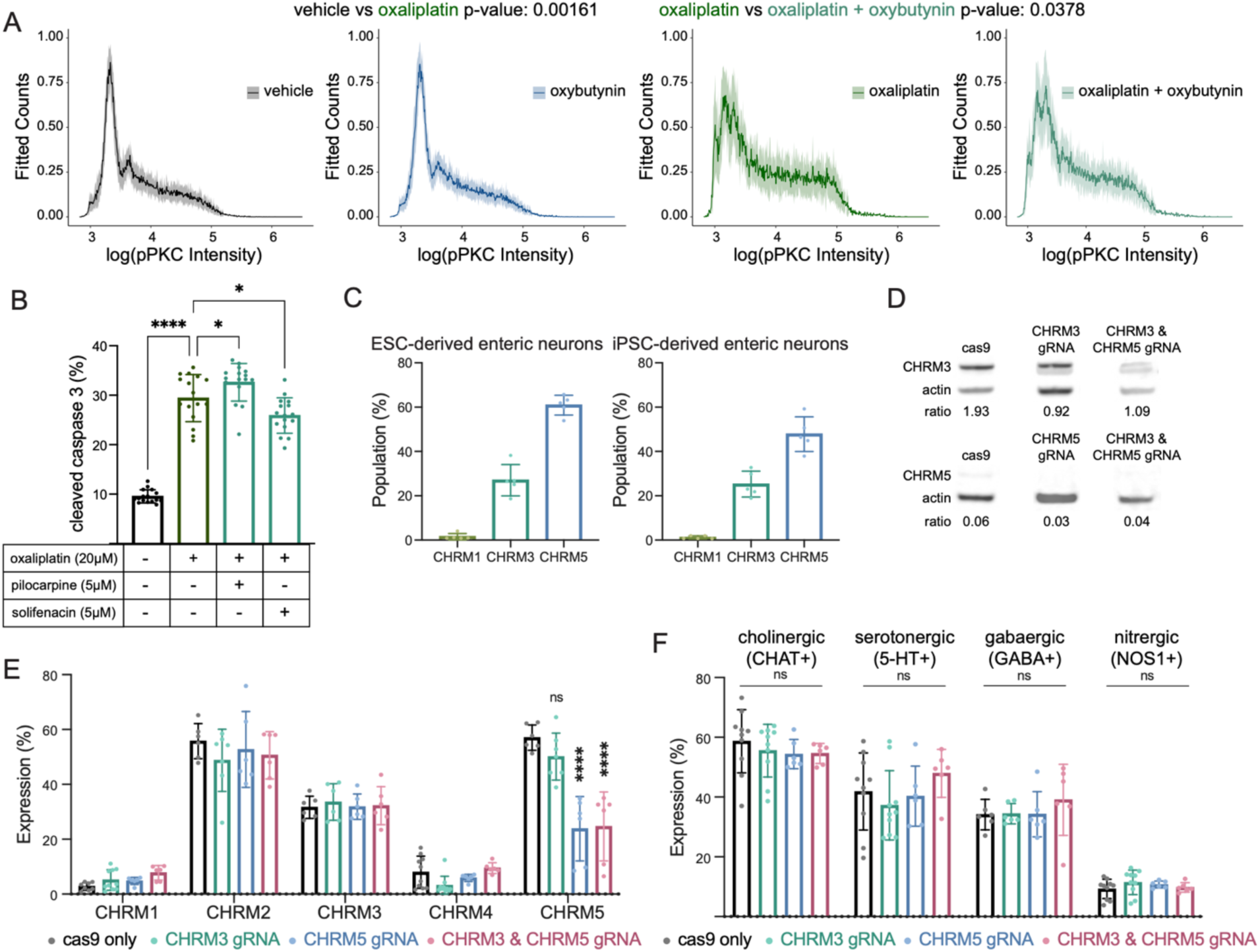
Effect of muscarinic antagonism on platin toxicity hallmarks and CHRM3 and CHRM5 CRISPR-cas9 targeted hPSC-derived enteric neuron characterization. **A)** Distribution of neuronal phosphorylated protein kinase C (pPKC) intensity per cell in 5 μM oxybutynin and 10 μM oxaliplatin-treated hPSC-derived enteric neurons. Vehicle contains water diluted to 1% in medium. Data are represented as mean ± SD. **B)** Quantification of flow cytometry analysis detecting cleaved caspase 3 levels in oxaliplatin-treated hPSC-derived enteric neurons co-treated with pilocarpine (muscarinic agonist) or solifenacin (muscarinic antagonist). Vehicle contains water diluted to 2% in medium. Data are represented as mean ± SD. **C)** Quantification of flow cytometry analysis detecting the expression of G_q_ muscarinic cholinergic receptors *CHRM1*, *CHRM3*, and *CHRM5* in ESC-derived and iPSC-derived enteric neurons. Data are represented as mean ± SD. **D)** Western blots and quantification of CHRM3 and CHRM5 protein levels in *CHRM3* and *CHRM5* CRISPR-cas9 RNP targeted hPSC-derived enteric neurons as compared to cas9 only targeted hPSC-derived enteric neurons. **E)** Quantification of flow cytometry analysis detecting the expression of muscarinic cholinergic receptors *CHRM1*, *CHRM2*, *CHRM3*, *CHRM4*, and *CHRM5* in *CHRM3* and *CHRM5* CRISPR-cas9 RNP targeted hPSC-derived enteric neurons as compared to cas9 only targeted hPSC-derived enteric neurons. Data are represented as mean ± SD. **F)** Quantification of flow cytometry analysis detecting the levels of neurotransmitter markers CHAT, GABA, NOS1, and serotonin in *CHRM3* and *CHRM5* CRISPR-cas9 RNP targeted hPSC-derived enteric neurons as compared to cas9 only targeted hPSC-derived enteric neurons. Data are represented as mean ± SD. *p value < 0.05, **** p value < 0.0001, ns: not significant.

**Supplementary Figure 4:**
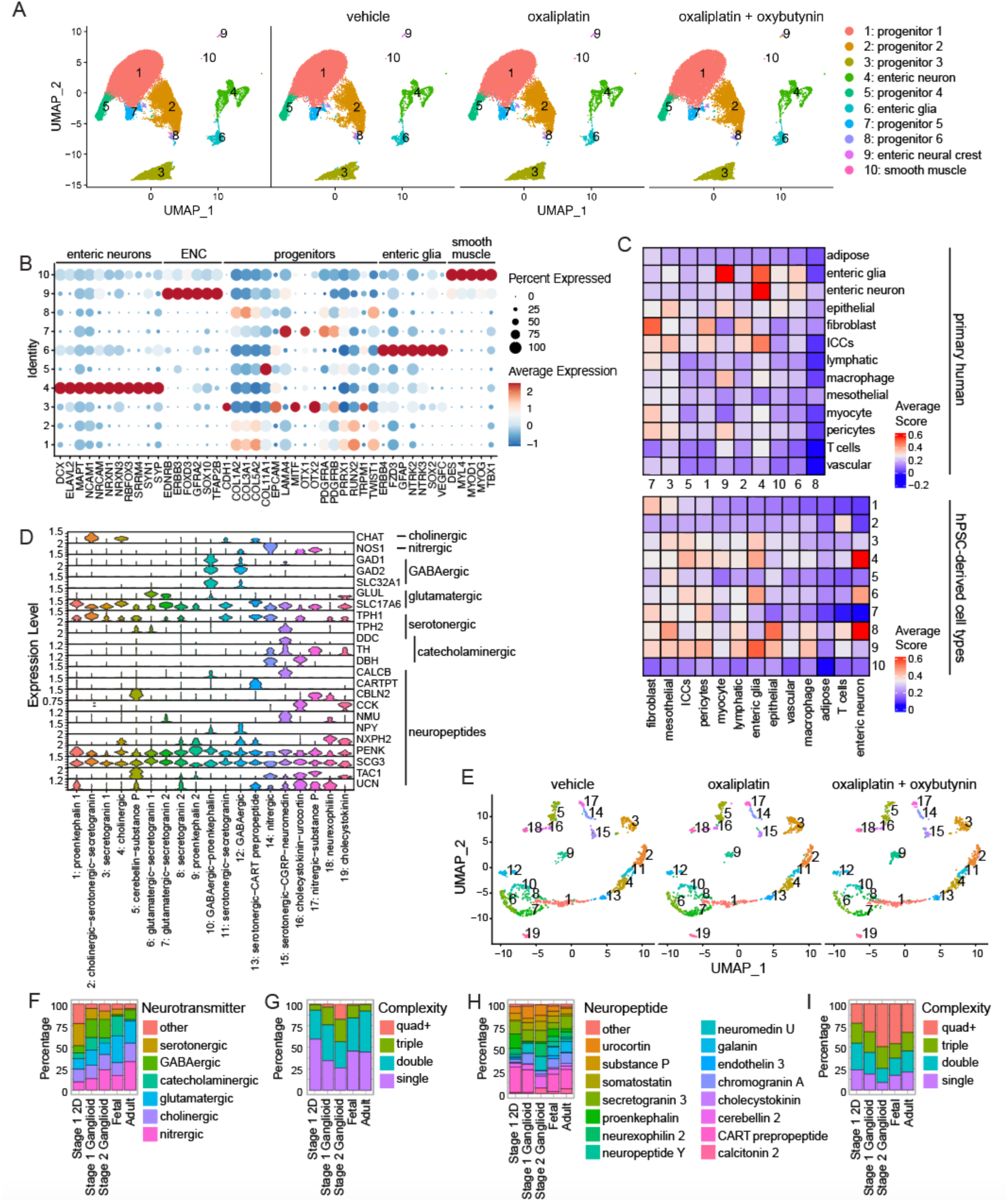
hPSC-derived 2D enteric neurons recapitulate human enteric neuron neurochemical diversity. **A)** sn-RNAseq UMAPs of cell types present in the hPSC-derived enteric neuron culture showing all cells (left) and the cells from each treatment condition: vehicle (medium containing 1% water), 10 μM oxaliplatin, and 10 μM oxaliplatin + 5 μM oxybutynin (right). **B)** Dot plot showing the expression of canonical enteric neuron, enteric neural crest (ENC), progenitor, enteric glia, and smooth muscle markers in the vehicle-treated stage 1 2D culture cell types. **C)** Heatmaps of the average module scores of adult human colon cell type transcriptional signatures in hPSC-derived cell types (top) and the average module scores of hPSC-derived cell type transcriptional signatures in adult human colon cell types (bottom). **D)** Dot plot showing the expression of enteric neuron neurochemical identity markers in the hPSC-derived enteric neuron subtypes. **E)** sn-RNAseq UMAPs of neuronal subtypes present in our hPSC-derived enteric neurons from each treatment condition: vehicle (medium containing 1% water), 10 μM oxaliplatin, and 10 μM oxaliplatin + 5 μM oxybutynin (right). **F-G)** Bar plots of the overall representation of neurotransmitter synthesizing neurons (left) and mono- and multi-neurotransmitter synthesizing neurons (right) in stage 1 2D enteric neurons compared to stage 1 and 2 enteric ganglioids and human primary fetal and adult enteric neurons. **H-I)** Bar plots of the overall representation of neuropeptide expressing neurons (left) and mono- and multi-neuropeptide expressing neurons (right) in stage 1 2D enteric neurons compared to stage 1 and 2 enteric ganglioids and human primary fetal and adult enteric neurons.

**Supplementary Figure 5:**
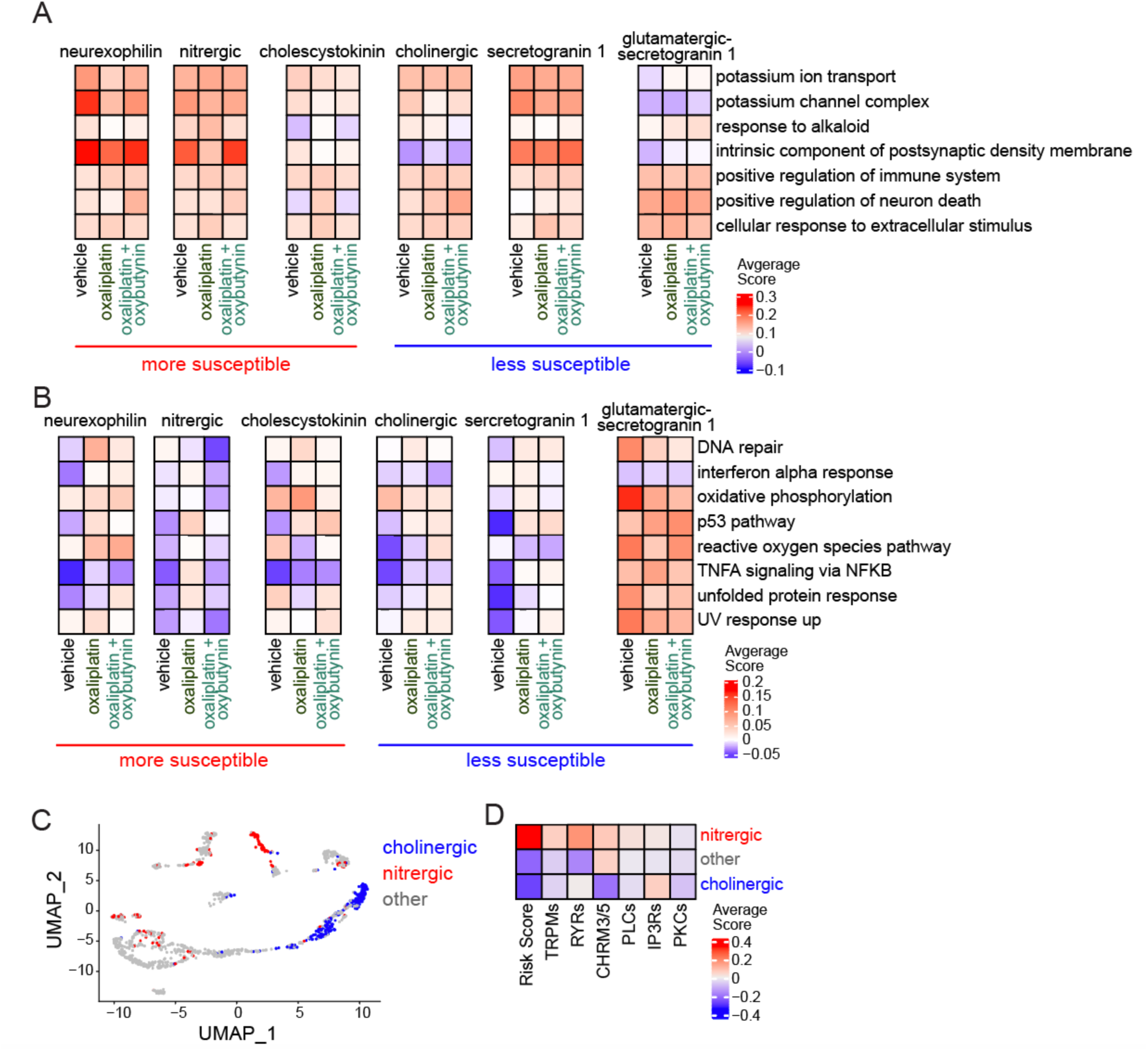
Cell type specific transcriptional profiling of platin-induced excitotoxicity susceptibility. **A)** Heatmaps of the average module scores of select Ontology neurotransmission and cellular stress related gene sets in the hPSC-derived enteric neuron subtypes predicted to be most susceptible (neurexophilin, nitrergic, and cholescystokinin) and least susceptible (cholinergic, secretogranin 1, and glutamatergic-secretogranin 1) to platin-induced excitotoxicity. **B)** Heatmaps of the average module scores of select Hallmark cellular stress related gene sets in the stage 1 2D enteric neuron subtypes predicted to be most susceptible (neurexophilin, nitrergic, and cholescystokinin) and least susceptible (cholinergic, secretogranin 1, and glutamatergic-secretogranin 1) to platin-induced excitotoxicity. **C)** sn-RNAseq UMAP of hPSC-derived enteric neurons predicted to be cholinergic (CHAT producing), nitrergic (NOS1 producing), or other (neither CHAT nor NOS1 producing). **D)** Heatmap comparing the average module scores of the platin excitotoxicity transcriptional program gene categories in the hPSC-derived enteric neurons predicted to be cholinergic (CHAT producing), nitrergic (NOS1 producing), or other (neither CHAT nor NOS1 producing).

## Supplementary Tables

**Table S1: Characteristics of individuals with colorectal cancer extracted from UCSF Research Data Browser**

**Table S2: Characteristics of individuals with ovarian cancer extracted from UCSF Research Data Browser**

**Table S3: Prevalence of constipation in patients that received platin chemotherapy and untreated patients**

**Table S4: Cytokine array results showing average MFI (n=3) normalized to vehicle**

**Table S5: List of predicted drug targets using compound-protein interaction analysis**

**Table S6: List of predicted targets for experimentally validated hit compounds**

**Table S7: List of transcripts and functional categories in excitotoxicity transcriptional program**

**Table S8: List of quality control metric cutoffs for the snRNAseq dataset**

**Table S9: Principal components used for SNN and UMAP calculation and the resolution used for clustering of each dataset**

**Table S10: List of transcripts used to define neurochemical identities**

**Table S11: List of antibodies**

## Methods

### Experimental Models Details

#### Human pluripotent stem cell lines

Human embryonic stem cell line H9 (WAe009-A, and derivatives *CHRM3* gRNA, *CHRM5* gRNA, cas9) and induced pluripotent stem cell line WTC-11 (UCSFi001-A) were plated on geltrex-coated plates and maintained in chemically-defined medium (E8) as described previously^23^. The maintenance cultures were tested for mycoplasma every 30 days.

#### Cancer cell lines

Colorectal cancer cell lines SW480 and WiDr were plated on geltrex-coated plates and maintained in chemically-defined mediums. SW480 was cultured in McCoy’s 5A medium (Corning, 10-050-CV) supplemented with 10% FBS, 0.5 mg/mL normocin (InvivoGen, ant-nr-2), and 1mM sodium pyruvate (Corning, 25-000-CI). WiDr was cultured in EMEM (VWR, 12-611F) supplemented with 0.5 mg/mL normocin (InvivoGen, ant-nr-2). Cultures were tested for mycoplasma every 30 days.

#### Mouse line

Male wild type C57BL/6J mice ages 8-12 weeks old (n = 40) were purchased from Jackson Laboratory. https://www.jax.org/strain/000664

### UCSF Research Data Browser Analysis

The UCSF Research Data Browser was utilized to identify and extract patients diagnosed with colorectal cancer (ICD-10-CM-C78.5) for the oxaliplatin analysis or ovarian cancer (ICD-10-CM-C56) for the carboplatin analysis. Patients diagnosed with constipation prior to their cancer diagnosis were excluded from both analyses, based on the ICD10 name “constipation”, to limit confounding factors. In each analysis, patients in the platin chemotherapy treated groups were then matched to their corresponding untreated groups by age, sex, and race/ethnicity. Patients were then further subset into four groups based on whether they were prescribed the relevant platin chemotherapy (based on the drug name (oxaliplatin or carboplatin)) and whether they were diagnosed with constipation (based on the ICD10 name “constipation”). Data represents the odds ratio and 95% confidence intervals for each drug calculated by Fisher’s exact statistical testing. p-values were calculated by Chi-squared statistical testing.

### Differentiation and Characterization of Enteric Neurons

#### Enteric neural crest induction

Once the hPSCs reached 70-80% confluency, the 12-day enteric neural crest (ENC) induction protocol was initiated according to our group’s established induction protocol^23^. Briefly, on day 0, the maintenance medium (E8) was aspirated and replacing with neural crest induction medium A [BMP4 (1 ng/mL), SB431542 (10 μM), and CHIR 99021 (600 nM) in Essential 6 medium]. Subsequently, on ENC induction days 2 and 4, the cells were fed with neural crest induction medium B [SB431542 (10 μM) and CHIR 99021 (1.5 μM) in Essential 6 medium] and on days 6, 8, and 10 the cells were fed with neural crest induction medium C [medium B with retinoic acid (1 μM)]. Next, enteric neural crestospheres were formed during days 12-15 in NC-C medium [FGF2 (10 ng/mL), CHIR 99021 (3 μM), N2 supplement (10 μL/mL), B27 supplement (20 μL/mL), glutagro (10 μL/mL), and MEM NEAAs (10 μL/mL) in neurobasal medium] following dissociation with Accutase. NC-C medium was refreshed on day 14 prior to enteric neuron induction phase on day 15.

#### Enteric neuron induction from enteric neural crest

On day 15 of the protocol, enteric neuron induction was initiated according to our group’s established induction protocol ^21,23^. From day 15 onward, cells were cultured in ENC medium [GDNF (10 ng/mL), ascorbic acid (100 μM), N2 supplement (10 μL/mL), B27 supplement (20 μL/mL), glutagro (10 μL/mL), and MEM NEAAs (10 μL/mL) in neurobasal medium] to promote the enteric neuron identity, with feeding occurring every two to three days. Enteric neuron progenitors (days 15-28) were dissociated with Accustase for 30 minutes 37°C, 5% CO2 and plated on poly-L-ornithine (PO)/laminin/fibronectin (FN) plates at 300,000 viable cells/cm^2^ in ENC medium and continued to be fed every two to three days until D30, after which, feeding frequency could be reduced to once or twice per week with larger volumes of feeding medium. All experiments were performed on stage 1 enteric neurons (days 35-50)^21^.

### Differentiation and Characterization of Control Peripheral Neurons

#### Cranial neural crest induction

Once the hPSCs reached 70-80% confluency, a previously established 12-day cranial neural crest induction protocol was initiated according to our group’s established induction protocol^46^. Briefly, on day 0, the maintenance medium (E8) was aspirated and replaced with neural crest induction medium A [BMP4 (1 ng/mL), SB431542 (10 μM), and CHIR 99021 (600 nM) in Essential 6 medium]. Subsequently, cells were fed with neural crest induction medium B [SB431542 (10 μM) and CHIR 99021 (1.5 μM) in Essential 6 medium] on days 2, 4, 6, 8, and 10. Next, cranial neural crest crestospheres were formed during days 12-15 in NC-C medium [FGF2 (10 ng/mL), CHIR 99021 (3 μM), N2 supplement (10 μL/mL), B27 supplement (20 μL/mL), glutagro (10 μL/mL), and MEM NEAAs (10 μL/mL) in neurobasal medium] following dissociation with Accutase (Stemcell Technologies, 07922). NC-C medium was refreshed on day 14 prior to the induction of cranial neural crest derived peripheral neurons on day 15.

#### Induction of cranial neural crest-derived peripheral neurons

On day 15, cranial neural crest-derived peripheral neuron induction was initiated according to our group’s established induction protocol^24,46^. From day 15 onward, cells were cultured in ENC medium [GDNF (10 ng/mL), ascorbic acid (100 μM), N2 supplement (10 μL/mL), B27 supplement (20 μL/mL), glutagro (10 μL/mL), and MEM NEAAs (10 μL/mL) in neurobasal medium] to promote the cranial neuron identities, with feeding occurring every two to three days. Cranial neural crest-derived neuron progenitors (days 15-28) were dissociated with Accustase (Stemcell Technologies, 07922) for 30 minutes 37°C, 5% CO2 and plated on poly-L-ornithine (PO)/laminin/fibronectin (FN) plates at 300,000 viable cells/cm^2^ in ENC medium and continued to be fed every two to three days until D30, after which, feeding frequency could be reduced to once or twice per week with larger volumes of feeding medium. All experiments were performed on cranial neural crest-derived neurons between D35 and D50.

### Immunofluorescence Staining of hPSC-derived Neurons

Cells were fixed in 4% paraformaldehyde (Santa Cruz, sc-281692) in PBS for 30 minutes at room temperature, and then blocked and permeabilized in eBioscience Foxp3/Transcription Factor Staining Buffer Set permeabilization buffer (Invitrogen, 00-5523) for another 30 minutes at room temperature. After fixation and permeabilization, cells were incubated in primary antibody solution diluted in permeabilization buffer overnight at 4°C, and then washed three times with permeabilization buffer before their incubation with fluorophore-conjugated secondary antibodies diluted in permeabilization buffer for one hour at room temperature. Before imaging, stained cells were incubated with DAPI fluorescent nuclear stain diluted in PBS and washed an additional two times with PBS. The list of antibodies and working dilutions is provided in Supplementary Table 11. Unless otherwise specified, cells were imaged on a digital inverted fluorescence microscope with a 20X objective and images were processed using Adobe Photoshop software.

### Preparation of Platins for *In Vitro* Experiments

Platins were prepared fresh for each experiment. Briefly, a stock solution of each platin was prepared by dissolving each compound in sterile water by vortexing vigorously. Stock solutions were 1 mM for cisplatin (Sigma-Aldrich, C2210000) and oxaliplatin (Sigma-Aldrich, Y0000271) and 10 mM for carboplatin (Sigma-Aldrich, C2538). Once in solution, each drug stock was diluted in cell culture medium to the designated concentration needed per experiment. The stock solutions were disposed of within 24 hours of preparation.

### Lactate Dehydrogenase (LDH) Activity Assay

To compare the cytotoxic response of the enteric neurons and control peripheral neurons to the platins, cells were assayed for lactate dehydrogenase activity using CytoTox 96 cytotoxicity assay kit (Promega, G1780). Briefly, the cells are plated in 96 well plates. The supernatant and the cell lysate are harvested three days later and assayed for lactate dehydrogenase activity using a plate reader (490 nm absorbance). Cytotoxicity is calculated by dividing the lactate dehydrogenase signal of the supernatant by the total lactate dehydrogenase signal (from lysate plus supernatant). Data were collected from nine independent differentiations. Statistical analysis was performed by two-way ANOVA with Šidák correction for multiple comparisons.

### Bulk RNA Sequencing Analysis

#### Sample preparation and sequencing

hPSC-derived enteric neurons and control peripheral neurons were treated with 50 µM oxaliplatin or vehicle (medium containing 5% water) for three days. Samples were prepared in duplicate, from two independent differentiations, per condition according to the QuantSeq 3′ mRNA-Seq Library Prep Kit FWD for Illumina (Lexogen). Libraries were sequenced on Hiseq 4000 at the UCSF Center for Advanced Technology.

#### Pre-ranked gene set enrichment analysis

Reads were aligned to the human genome (build hg38) using STAR (v2.7.1a). FeatureCounts (v1.6.2) were then used to count reads assigned to gene features, as annotated in Illumina’s iGenome hg38 GTF (2015). Gene level counts were then assessed for differential expression analysis with DESeq2 (v1.20) in the R (v3.5.1) environment. Briefly, the DESeq function was used with default options for each comparison: oxaliplatin-treated control peripheral neurons vs vehicle-treated control peripheral neurons, oxaliplatin-treated enteric neurons vs vehicle-treated enteric neurons, and oxaliplatin-treated enteric neurons vs oxaliplatin-treated control peripheral neurons. The differentially expressed genes were sorted in descending order by the log_2_ fold-change in gene expression for each comparison. Then pre-ranked GSEA was performed using GSEA version 4.3.2 and the MSigDB Hallmark and Ontology gene sets version 2022.1 to identify gene sets significantly changed between the two populations based on a p-value cutoff of < 0.05 and a false discovery rate of < 0.25.

### Cytokine ELISA

#### Sample preparation

hPSC-derived enteric neurons and hPSC-derived control peripheral neurons cultured in 24-well plates were treated with 20 µM cisplatin, 200 µM carboplatin, 40 µM oxaliplatin, or vehicle (medium containing 4% water) for three days. Three biological replicates per treatment condition were included, for a total of 24 samples. Cell culture supernatants were transferred to 1.5 mL tubes and centrifuged at 4°C and 10,000g for 10 minutes. Each supernatant was transferred to fresh 1.5 mL tubes, frozen at −80°C, and shipped overnight on dry ice to the Human Immune Monitoring Center at Stanford University.

#### Luminex assay

This assay was performed by the Human Immune Monitoring Center at Stanford University-Immunoassay Team. Kits were purchased from EMD Millipore Corporation, Burlington, MA., and run according to the manufacturer’s recommendations with modifications described as follows. H76 kits include 3 panels: Panel 1 is Milliplex HCYTMAG-60K-PX41 with addition of IL-18 and IL-22. Panel 2 is Milliplex HCP2MAG-62K-PX23 with addition of MIG. Panel 3 includes the Milliplex HSP1MAG-63K-06 and HADCYMAG-61K-03 (Resistin, Leptin and HGF) to generate a 9 plex. Samples were diluted 3-fold (Panels 1 and 2) and 10-fold for Panel 3. An aliquot (25 µL) of the diluted sample was mixed with antibody-linked magnetic beads in a 96-well plate and incubated overnight at 4°C with shaking. Cold and room temperature incubation steps were performed on an orbital shaker at 500-600 rpm. Plates were washed twice with wash buffer in a BioTek ELx405 washer (BioTek Instruments). Following one-hour incubation at room temperature with biotinylated detection antibody, streptavidin-PE was added for 30 minutes with shaking. Plates were washed as described above and PBS added to wells for reading in the Luminex FlexMap3D Instrument with a lower bound of 50 beads per sample per cytokine. Each sample was measured in duplicate. Custom Assay Chex control beads were purchased and added to all wells (Radix BioSolutions). Wells with a bead count <50 were flagged, and data with a bead count <20 were excluded.

#### Report and data analysis

The output CSV file was used to extract the median fluorescence intensities for each cytokine. The median fluorescence intensities per sample were averaged. Then the average median fluorescence intensities for each biological replicate were normalized to the average of each population’s vehicle treatment condition. The average of the three biological replicates for each cytokine are represented in the heatmaps. Statistical analysis was performed by two-way ANOVA with Dunnett correction for multiple comparisons.

### Viability Assay

To compare the cell viability of the enteric neurons treated with the platins with and without a variety of chemical modulators, cells were assayed for ATP levels using CellTiter-Glo Luminescent Cell Viability assay kit (Promega, G7571). Briefly, cells were plated in 96 well plates and treated with the experiment treatment conditions, specified below, for three days and then assayed for ATP levels using a plate reader, detecting luminescence with a 500 ns integration time. The ATP levels per well were then normalized to the average value of the vehicle wells to determine the normalized cell viability.

#### Platin cell viability experiment

Cells were treated with cisplatin at 1.25, 2.5, 5, and 10 µM concentrations, carboplatin at 20, 40, 80, and 100 µM, and oxaliplatin at 2.5, 5, 10 and 20 µM concentrations. Controls included 20 µM lomustine, 10 µM CCCP, and vehicle containing medium with 2% water. Data represents the mean ± SEM from six individually treated wells per condition. Statistical analysis was performed by two-way ANOVA with Dunnett correction for multiple comparisons.

#### High-throughput phenotypic screen compound validation experiment

Oxaliplatin was dosed at 40 µM. Each compound (mefloquine, mitotane, and oxybutynin) was dosed at 1 and 10 µM concentrations. Vehicle contained medium with 4% water. Each dot represents an individually treated well. Statistical analysis was performed by one-way ANOVA with Tukey correction for multiple comparisons.

#### G_q_ protein experiment

Oxaliplatin was dosed at 20 µM. YM-24489 was dosed at 2.5, 5, and 10 µM concentrations. Vehicle contained medium with 2% water. Each dot represents an individually treated well. Statistical analysis was performed by one-way ANOVA with Tukey correction for multiple comparisons.

#### G_i_ protein experiment

Oxaliplatin was dosed at 20 µM. Pertussis toxin was dosed at 0.01, 0.1, and 0.5 µg/mL concentrations. Vehicle contained medium with 2% water. Each dot represents an individually treated well. Statistical analysis was performed by one-way ANOVA with Tukey correction for multiple comparisons.

#### Phospholipase C experiment

Oxaliplatin was dosed at 20 µM. U 73122 was dosed at 2.5, 5, and 10 µM concentrations. Vehicle contained medium 2% water. Each dot represents an individually treated well. Statistical analysis was performed by one-way ANOVA with Tukey correction for multiple comparisons.

#### IP3 receptor experiment

Oxaliplatin was dosed at 20 µM. Lanthanum(III) was dosed at 100, 200, and 400 µM concentrations. Vehicle contained medium with 2% water. Each dot represents an individually treated well. Statistical analysis was performed by one-way ANOVA with Tukey correction for multiple comparisons.

#### Protein kinase C experiment

Oxaliplatin was dosed at 20 µM. Go 6983 was dosed at 0.1, 1, and 10 µM concentrations. Vehicle contained medium with 2% water. Each dot represents an individually treated well. Statistical analysis was performed by one-way ANOVA with Tukey correction for multiple comparisons.

### γH2AX Imaging and Analysis

Cells were plated in 24 well plates and treated with 5 and 10 µM cisplatin, 50 and 100 µM carboplatin, 10 and 20 µM oxaliplatin, or vehicle containing medium with 2% water for three days. Cells were fixed and stained with γH2AX and TUBB3 antibodies and DAPI, as described previously. High-throughput imaging was carried out using the In Cell Analyzer 2000 with a 20X objective (GE Healthcare). Neuronal γH2AX signal was detected based on the overlap of γH2AX with TUBB3. The total number of neuronal γH2AX pixels were normalized by the total cell number within each well as measured by DAPI staining with the GE Developer Toolbox v1.9.1. The quantification per well was then normalized to the average value of the vehicle wells. Each dot represents an individually treated well. Statistical analysis was performed by one-way ANOVA with Dunnett correction for multiple comparisons. Representative images were taken on the EVOS^TM^ FL digital inverted fluorescence microscope (Invitrogen).

### mitoSOX Imaging and Analysis

Cells were plated in 24 well plates and treated with 5 and 10 µM cisplatin, 50 and 100 µM carboplatin, 10 and 20 µM oxaliplatin, or vehicle containing medium with 2% water for three days. Cells were incubated in 5 µM mitoSOX reagent (Invitrogen, M36008) for 10 minutes at 37°C, then fixed and stained with TUBB3 and DAPI, as described previously. High-throughput imaging was carried out using the In Cell Analyzer 2000 with a 20X objective (GE Healthcare). Neuronal mitoSOX signal was detected based on the overlap of mitoSOX with TUBB3. The total number of neuronal mitoSOX pixels were normalized by the total cell number within each well as measured by DAPI staining with the GE Developer Toolbox v1.9.1. The quantification per well was then normalized to the average value of the vehicle wells. Each dot represents an individually treated well. Statistical analysis was performed by one-way ANOVA with Dunnett correction for multiple comparisons. Representative images were taken on the EVOS^TM^ FL digital inverted fluorescence microscope (Invitrogen).

### Neurite Length Imaging and Analysis

Cells were plated in 24 well plates and treated with 5 and 10 µM cisplatin, 50 and 100 µM carboplatin, 10 and 20 µM oxaliplatin, or vehicle containing medium with 2% water for three days. Cells were stained with TUBB3 and DAPI, as described previously. High-throughput imaging was carried out using the In Cell Analyzer 2000 with a 20x objective (GE Healthcare). Images were batch processed through an imaging processing software, MIPAR Image Analysis, with a custom-built algorithm to analyze measurements for chemotherapy-induced neurite degeneration, which has been published previously^29,30^.

Briefly, the algorithm generates optimized grayscale images by reducing overall noise and minimizing the amount of nonspecific staining to identify and quantify the neurite networks within each field-of-view image. A subsequent segmentation algorithm was performed to identify and quantify nuclei within each field-of-view image. After processing, each image yielded measurements of total neurite length and cell number. The total length of neurites per well were normalized by the total cell number per well. Each dot represents an individually treated well. Statistical analysis was performed by one-way ANOVA with Dunnett correction for multiple comparisons. The quantification per well was then normalized to the average value of the vehicle wells.

### Flow cytometry

Cells were dissociated into single cell suspensions by Accutase treatment (Stemcell Technologies, 07922) for 30-60 min, 37°C, 5% CO_2_ and then fixed and permeabilized using the eBioscience Foxp3/Transcription Factor Staining Buffer Set (Invitrogen, 00-5523). Cells were stained with primary and secondary antibodies as described previously for immunofluorescence. Flow cytometry was conducted using a flow cytometer and data was analyzed using FlowjoTM (FlowJoTM Software Version 8.7). The list of antibodies and working dilutions is provided in Supplementary Table 11.

### Cleaved Caspase 3 Detection and Analysis

#### Flow cytometry

Cells were plated in either 96 well or 24 well plates and treated with the experiment treatment conditions, specified below, for three days. Cells were stained with cleaved caspase 3 and TUBB3 antibodies and flow cytometry was performed, as described above. In FlowjoTM, neurons were detected by gating on the TUBB3+ cells. Then, within the TUBB3+ population, the percentage of neurons positive for cleaved caspase 3 was detected and reported.

#### Platin apoptosis experiment

Cisplatin was dosed at 5 and 10 µM. Carboplatin was dosed at 50 and 100 µM. Oxaliplatin was dosed at 10 and 20 µM. The vehicle condition contained medium with 2% water. Each dot represents an individually treated well. Statistical analysis was performed by one-way ANOVA with Dunnett correction for multiple comparisons.

#### Muscarinic cholinergic receptor agonist/antagonist experiment

Oxaliplatin was dosed at 20 µM. Pilocarpine and solifenacin were dosed at 5 µM. The vehicle condition contained medium with 2% water. Each dot represents an individually treated well. Statistical analysis was performed by one-way ANOVA with Dunnett correction for multiple comparisons.

#### CHRM3 and CHRM5 CRISPR RNP experiment

Oxaliplatin was dosed at 10 µM. Oxybutynin was dosed at 5 µM. The vehicle condition contained medium with 1% water. All targeted cell lines were treated, fixed, and stained for flow cytometry on the same days. Each dot represents an individually treated well. Statistical analysis was performed by two-way ANOVA with Tukey correction for multiple comparisons.

#### Four enteric neurotransmitter identities experiment

Oxaliplatin was dosed at 20 µM and the vehicle condition contained medium with 2% water. Each cleaved caspase 3 and TUBB3 antibody panel included one of four enteric neurotransmitter markers: NOS1, GABA, CHAT, and 5-HT. The list of antibodies and working dilutions is provided in Supplementary Table 11. Each dot represents an individually treated well. Statistical analysis was performed by two-way ANOVA with Šidák correction for multiple comparisons.

#### Imaging

Cells were plated in 96 well plates and treated with 10 µM oxaliplatin with or without oxybutynin dosed at 0.1, 1 and 5 µM, or vehicle containing medium with 1% water for two days. Cells were stained with cleaved caspase 3 and TUBB3 antibodies and stained with DAPI, as described previously. High-throughput imaging was carried out using the ImageXpress Confocal HT.ai (Molecular Devices). MetaXpress High-Content Image Acquisition and Analysis Software. Representative images were taken on the Revolve digital fluorescence microscope (ECHO). Each dot represents an individually treated well. Statistical analysis was performed by one-way ANOVA with Tukey correction for multiple comparisons.

### High-Throughput Phenotypic Drug Screen

Cells were plated in 384 well plates and treated with 40 µM oxaliplatin and compounds from an FDA-approved chemical library (Selleckchem, USA) at 1 µM. Three days after treatment, cells were stained with DAPI prior to fixation. Cells were then promptly fixed, as described previously, and stained with propidium iodide. High-throughput imaging was carried out using the In Cell Analyzer 2000 (GE Healthcare). Cell viability was quantified by using the propidium iodide signal to count the total number of cells per well and using the DAPI signal to count the total number of dead cells per well in the GE Developer Toolbox v1.9.1. Cell viability z-scores were calculated for each drug by subtracting the mean of the library from the drug and dividing that by the standard deviation of the library. This experiment was performed in two biological replicates, in ESC-derived enteric neurons and iPSC-derived enteric neurons. The top 30 drugs with the highest cell viability z-scores from each biological replicate were defined as hits from the screen.

### Colorectal Cancer Viability Imaging and Analysis

One day after passaging, cancer cells were treated with 20 µM cisplatin, 200 µM carboplatin, and 20 µM oxaliplatin and mefloquine, mitotane, and oxybutynin at 1, 4, 7, and 10 µM concentrations. Three days after treatment, cells were stained with DAPI prior to fixation. Cells were then promptly fixed, as described previously, and stained with propidium iodide. High-throughput imaging was carried out using the In Cell Analyzer 2000 (GE Healthcare). Cell viability was quantified by using the propidium iodide signal to count the total number of cells per well and using the DAPI signal to count the total number of dead cells per well in the GE Developer Toolbox v1.9.1. Each dot represents an individually treated well. Statistical analysis was performed by one-way ANOVA with Dunnett correction for multiple comparisons.

### Drug-Protein Interaction Pipeline

For this analysis, we used our previously published drug-protein interaction analysis pipeline to predict drug targets enriched in the compounds that either increase or decrease oxaliplatin-treated enteric neuron viability^32^. Briefly, isomeric SMILES for each drug in the library were acquired from Selleckchem and used to run a SEA library search. The SEA predicted targets were filtered, selecting human targets and predicted interaction *p*-values < 0.05, which yielded 2150 predicted proteins targeted by the drug library. Weighted combined z-scores were then calculated for each gene by combining *z*-scores across all treatments. The *p*-values were then calculated based on the combined z-scores and adjusted using p.adjust (method = FDR). As an orthogonal approach for each gene, we recorded the number of treatments with negative or positive *z*-scores as well as the total number of compounds predicted to target that gene. Using the sum of counts for all other genes and drugs, we performed a Fisher’s exact test to evaluate the degree to which *z*-scores were enriched among the treatments that either increase or decrease oxaliplatin-treated enteric neuron viability. A false discovery rate < 0.25 and Fisher’s p-value < 0.05 was used to identify the proteins significantly associated with a high or low viability z-score.

### Protein-Protein Interaction Network Analysis

Protein-protein interaction network analysis was performed using the Search Tool for the Retrieval of Interacting Genes (STRING) database version 11.0. The minimum required interaction score was set to 0.7, corresponding to high confidence with data support from the following active interaction sources: textmining, experiments, databases, co-expression, neighborhood, gene fusion, and co-occurrence.

### Microelectrode Array Analysis

#### Data acquisition

Cells were plated on Cytoview MEA 6-well plates and treated with 10 µM oxaliplatin or vehicle containing medium with 1% water. For the experiment including oxybutynin, enteric neurons were treated with 10 µM oxaliplatin, 5 µM oxybutynin, 10 µM oxaliplatin and 5 µM oxybutynin co-treatment, or vehicle containing medium with 1% water. After three hours, enteric neuron activity was recorded with the Axion Maestro Edge for 1 hour.

#### Data processing

Raw data were first spike sorted with a modified version of SpikeInterface (https://github.com/SpikeInterface) using MountainSort to identify high quality units by manually scoring based on amplitude, waveform shape, firing rate, and inter-spike interval contamination. Units with firing rates greater than 0.1 Hz were included in the analysis. Each dot represents an individual unit identified from spike sorting that passed scoring. Statistical analysis was performed by Welch’s t test for the experiment comparing oxaliplatin to vehicle and Brown-Forsythe and Welch ANOVA with Dunnett correction for multiple comparisons for the experiment including oxybutynin.

### Terminal Deoxynucleotidyl Transferase dUTP Nick End Labeling (TUNEL) Imaging and Analysis

TUNEL staining was performed using the In Situ Cell Death Detection Kit, TMP Red kit (Roche, 12156792910). Briefly, cells were plated in 96 well plates and treated with 10 µM oxaliplatin with or without oxybutynin dosed at 0.1, 1 and 5 µM, or vehicle containing medium with 1% water for three days. Cells were stained with TUBB3, as described previously, then stained with TUNEL enzyme solution diluted 10-fold in label solution by incubating each well in 50 µL of the diluted enzyme solution for sixty minutes at 37°C. Cells were washed twice with PBS then stained with DAPI as described previously. High-throughput imaging was carried out using the ImageXpress Confocal HT.ai (Molecular Devices). Neuronal TUNEL signal was detected based on the overlap of TUNEL with TUBB3. Neuronal TUNEL intensity was normalized by the total cell number within each well as measured by DAPI staining with the MetaXpress High-Content Image Acquisition and Analysis Software. Each dot represents an individually treated well. Statistical analysis was performed by one-way ANOVA with Tukey correction for multiple comparisons.

### Phosphorylated Protein Kinase C Imaging and Analysis

Cells were plated in 96 well plates and treated with 10 µM oxaliplatin with or without oxybutynin dosed at 5 µM, or vehicle containing medium with 1% water for three days. Cells were stained with p-PKC and TUBB3 antibodies then stained with DAPI as described previously. High-throughput imaging was carried out using the ImageXpress Confocal HT.ai (Molecular Devices). Neuronal p-PKC signal was detected based on the overlap of p-PKC with TUBB3. Neuronal p-PKC intensity was normalized by the total cell number within each well as measured by DAPI staining with the MetaXpress High-Content Image Acquisition and Analysis Software. Each dot represents an individually treated well. Statistical analysis was performed by one-way ANOVA with Holm-Šidák correction for multiple comparisons. The distribution of neuronal p-PKC intensity per cell was also visualized by binning log_10_ transformed intensity values across 512 equally sized bins between the minimum and maximum intensity value among all treatment condition wells. Distributions were scaled per well. Distribution plots were generated using ggplot2 (v3.4). The solid line represents the average distribution of all wells from the same treatment condition with the standard deviation shown by the transparent ribbon. Statistical analysis was performed by nested t-tests comparing the following conditions: vehicle vs oxaliplatin and oxaliplatin vs oxaliplatin + oxybutynin. Representative images were taken on the Revolve digital fluorescence microscope (ECHO) and processed using Fiji (ImageJ) software.

### CHRM3 and CHRM5 CRISPR Ribonucleoprotein (RNP) Targeting

#### Targeting

*CHRM3* and *CHRM5* ribonucleoprotein (RNP) complexes were assembled by mixing 16 pmol of multi-guide sgRNA (Synthego) and 12 pmol of Cas9 2NLS (Synthego) in lipofectamine CRISPRMAX Cas9 transfection reagent (Invitrogen, CMAX00001) per reaction. H9 ESCs were lipofected with the RNP-transfection solutions then replated after two days in 10 cm dishes at 1500 cells/cm^2^. After a few days, individual colonies were picked and plated into individual wells of a 24 well plate. After a few more days, the clones were frozen at −80°C in STEM-CELLBANKER DMSO FREE (Amsbio LCC, 13926) and stored long-term in liquid nitrogen.

#### Protein extraction

hPSC-derived enteric neurons were lysed in RIPA lysis buffer (Millipore, 20-188) with protease/phosphatase inhibitor (Roche, 4693159001) at 4°C for 30 minutes while rocking. Lysate was centrifuged at 13,800 g for 20 minutes at 4°C. Supernatants were transferred to fresh tubes and stored long-term at −80°C.

#### Protein quantification

Bovine serum albumin (BSA, Sigma-Aldrich, A4503) was diluted to 1, 0.8, 0.6, 0.4, 0.2, 0.1, and 0.05 mg/mL with water. Lysates were diluted 1:1, 1:10, and 1:100 in water. Lysates and BSA samples were added to Bio-Rad Protein Assay Dye Reagent Concentrate (Bio-Rad Laboratories, 5000006) and incubated for 10 minutes before absorbance was measured at 595 nm on a plate reader. BSA concentrations versus the BSA absorbances were used to generate a standard curve and calculate the lysate protein concentrations.

#### Western blot

Denaturing protein gel electrophoresis was performed according to Invitrogen NuPAGE Bis-Tris Mini Gels user guide. Briefly, protein samples were denatured in LDS Buffer (Invitrogen, NP0007) and Reducing Agent (Invitrogen, NP0009) for 10 minutes at 70°C. Each sample (15 µg) was loaded and run in a NuPAGE 4-12% Bis-Tris gel (Invitrogen, NP0322BOX) on the Mini Gel Tank (Invitrogen) for 45 minutes at 150 volts. Western blotting was then performed according to the Invitrogen General Procedure for Chemiluminescent Western Blotting. Briefly, samples were transferred using the Power Blotter Station (Invitrogen) for seven minutes at 25 volts and 2.5 amps. After transfer, the membrane was blocked in 2% milk and 2% BSA in tris buffered saline with 0.05% tween 20 for one hour with gentle rocking. The membrane was then incubated in CHRM3, CHRM5, and actin antibodies diluted in 2% milk and 2% BSA in tris buffered saline with 0.05% tween 20 at the designated concentrations overnight at 4°C with gentle rocking. The list of antibodies and working dilutions is provided in Table 2.11. The membrane was then washed three times in tris buffered saline with 0.05% tween 20 for 10 minutes each wash with gentle rocking. The membrane was then incubated in HRP conjugated secondary antibodies diluted in 2% milk and 2% BSA in tris buffered saline with 0.05% tween 20 for two hours with gentle rocking. The membrane was washed again as described above, developed in chemiluminescent substrate, and then immediately imaged on the Odyssey Fc (LI-COR). Protein bands were quantified in Fiji (ImageJ).

### Single Nuclei RNA Sequencing

#### Sample preparation and data collection

Cells were plated in 24 well plates and treated with 10 µM oxaliplatin, 10 µM oxaliplatin with 5 µM oxybutynin, and vehicle containing medium with 1% water for 24 hours. Cells were dissociated into single cell suspensions by Accutase treatment (Stemcell Technologies, 07920) for 30 minutes at 37 °C, 5% CO_2_. Single nuclei suspensions were prepared in duplicate per condition according to 10x Genomics Isolation of Nuclei for Single Cell Sequencing Demonstrated Protocol. Briefly, dissociated cells were lysed on ice in lysis buffer for two minutes and washed with nuclei wash and resuspension buffer and resuspended to 1000 nuclei/µL in nuclei wash and resuspension buffer. Cells were strained to remove cell debris and large clumps then immediately submitted to the UCSF Genomics CoLab for GEM generation & barcoding, post GEM-RT cleanup & cDNA amplification, 3’ gene expression library construction, and sequencing on Illumina NovaSeq sequencer according to the 10x Genomics Chromium Next GEM Single Cell 3’ Reagent Kits v3.1 User Guide. Gene count matrices were generated using CellRange (v6.0) with alignment to the human reference hg38 transcriptome.

#### Quality control and cell filtration

Datasets were analyzed in R version 4.2.1 with Seurat version 4.3.0^47^. The number of reads mapping to mitochondrial and ribosomal gene transcripts per cell were calculated using the “PercentageFeatureSet” function. Cells were identified as poor quality and subsequently removed independently for each dataset based on the number of unique features captured per cell, the number of UMI captured per cell and the percentage of reads mapping to mitochondrial transcripts per cell. Dataset specific quality control metric cutoffs can be found in Supplementary Table 8.

#### Dimensionality reduction, clustering, and annotation

For each biological replicate, quality control metrics were visualized to identify and remove low quality cells (Supplementary Table 8). Counts matrices were log normalized with a scaling factor of 10,000 and 2,000 variable features were identified using the “vst” method. Count matrices were integrated using Seurat integration functions with default parameters. The variable feature sets were scaled and centered. Principal Components Analysis (PCA) was run using default settings and Uniform Manifold Approximation and Projection (UMAP) dimensionality reduction was performed using the PCA reduction. The shared nearest neighbors (SNN) graph was computed using default settings and cell clustering was performed using the default Louvain algorithm. The number of principal components used for UMAP reduction and SNN calculation was determined by principal component standard deviation and varied for each dataset and can be found in Supplementary Table 9. The above pipeline was performed again for sub-clustering enteric neurons from each biological replicate. Biological replicates per treatment condition were merged using the base R “merge” function and the above pipeline was repeated for each treatment condition. Gene dropout values were imputed using adaptively-thresholded low rank approximation (ALRA) for each treatment condition^48^. The rank-k approximation was automatically chosen for each dataset and all other parameters were set as the default values. The imputed gene expression was used to identify cluster-specific expression of cell type marker genes and neurochemical identity genes for the all cells dataset and sub-clustered enteric neuron dataset, respectively.

#### Curation of published datasets

Curation of the snRNA-seq datasets of primary adult human colon and hPSC-derived stage 1 and stage 2 ganglioid cultures was conducted as previously described by our group ^21^. For the primary fetal human enteric neuron single cell RNA sequencing dataset, previously published by Teichmann and colleagues, the normalized .H5AD file was downloaded from https://www.gutcellatlas.org/ and converted into a Seurat object.

#### Cell type transcriptional signature scoring

To find transcriptionally similar cell populations between our dataset and the primary human adult GI tissue dataset, the differentially expressed genes of the reference dataset were first calculated from the non-imputed gene counts with the “FindAllMarkers” function using the Wilcoxon Rank Sum test and only genes with a positive fold change were returned. Since it was unclear how the drug treatments might affect the genes that are differentially expressed, this analysis was completed with the vehicle-treated cells from our dataset. The query dataset was then scored for the transcriptional signature of each reference dataset cell cluster using the “AddModuleScore” function based on the query dataset’s imputed gene counts.

#### Neurochemical identification of neurons

The neurochemical identification of neurons was performed independently for each neurotransmitter and neuropeptide to accommodate multi-neurochemical identities. Since it was unclear how the drug treatments might affect the expression of neurochemical identity genes, this analysis was completed with the vehicle-treated neurons from our dataset. For each neurotransmitter, a core set of genes were selected consisting of the rate-limiting synthesis enzyme(s), metabolism enzymes and transport proteins (Supplementary Table 10). Cells were first scored for each neurotransmission associated gene set using the “AddModuleScore” function. A cell was then annotated as “x-ergic” if the cell’s expression of a rate limiting enzyme was greater than 0 and the cell’s module score for the corresponding gene set was greater than 0. A cell was annotated as “other” if both criteria were not met. For each neuropeptide, cells were scored for the gene corresponding with each neuropeptide using the “AddModuleScore” function (Supplementary Table 10). A cell was then annotated as the neuropeptide if the cell’s expression of a gene was greater than 0 or “other” if less than 0. Multi-neurochemical identities were determined by concatenating the individually determined single neurochemical identities of each cell. The overall prevalence of each neurochemical identity per dataset was calculated by summing the total number of cells annotated for each single identity and calculating the percentage of each identity from this sum total.

#### Excitotoxicity transcriptional program scoring

To identify the expression level of the excitotoxicity transcriptional program genes in the enteric neuron populations, cells were scored for their expression of each platin excitotoxicity transcriptional program gene category as well as a combined list of all genes in the platin excitotoxicity transcriptional program using the “AddModuleScore” function (Supplementary Table 7).

#### Pre-ranked gene set enrichment analysis

Genes differentially expressed between the vehicle and oxaliplatin treatment conditions and the oxaliplatin treatment and oxaliplatin-oxybutynin co-treatment conditions for the neurexophilin, nitrergic, cholecystokinin, cholinergic, secretogranin 1 and glutamatergic-secretogranin 1 populations were identified using the “FindAllMarkers” function using the Wilcoxon Rank Sum test. Pre-ranked GSEA for the MSigDB Hallmark and Ontology gene sets was performed on each subtypes sorted by decreasing log_2_ fold-change using fgsea v1.16. Normalized enrichment scores were calculated for gene sets containing a minimum of 15 or maximum of 500 genes in the differentially expressed gene list. Gene sets significantly changed between the two treatment conditions were identified based on a p-value cutoff of < 0.05. Gene sets rescued by oxybutynin treatment were identified based on the gene set’s normalized enrichment score being reversed (positive to negative or negative to positive) when comparing the vehicle versus oxaliplatin pre-ranked GSEA results to the oxaliplatin versus oxaliplatin-oxybutynin pre-ranked GSEA results.

#### MSigDB gene set scoring

Separate gene lists were created containing all genes belonging to each MSigDB Hallmark or Ontology gene set of interest. Cells were scored for their expression of each gene list using the “AddModuleScore” function. For the MSigDB Hallmark gene sets, a combined list of all genes in the Hallmark gene sets was used to generate a combined cellular stress score.

### Nitric Oxide Release Assay

Cells were plated in 96 well plates. Pre-treatment supernatants were collected by washing the cells with Tyrode’s solution [NaCl (129 mM), KCl (5 mM), CaCl_2_ (2 mM), MgCl_2_ (1 mM), glucose (30 mM) and HEPES (25 mM) at pH 7.4]. Cells were then incubated with Tyrode’s solution or 50 µM acetylcholine diluted in Tyrode’s solution for 1 hour at 37°C, 5% CO_2_. Supernatants were then removed, centrifuged at 2313g for 10 minutes at 4°C, and frozen at −80°C. Cells were then treated with 10 µM oxaliplatin, vehicle containing medium with 1% water, or 10 µM CCCP for 24 hours. Cells were washed with Tyrode’s solution and incubated with Tyrode’s solution or 50 µM acetylcholine diluted in Tyrode’s solution for 1 hour at 37°C, 5% CO_2_. Supernatants were removed, centrifuged at 2313g for 10 minutes at 4°C, and frozen at −80°C. Nitric oxide levels in the supernatants were determined using a nitric oxide assay kit (Invitrogen, EMSNO). Briefly, the kit uses the enzyme nitrate reductase to convert nitrate to nitrite which is then detected as a colored azo dye absorbing light at 540 nm. Nitric oxide release for each well was normalized to the average absorbance of the pre-treatment no acetylcholine stimulation wells. Each dot represents an individually treated well. Statistical analysis for the t=0 hour time point was performed by Welch’s t test. Statistical analysis for the t=24 hour time point was performed by two-way ANOVA with Dunnett correction for multiple comparisons.

### Preparation of Paraffin-Embedded Human Stomach Sections

#### Tissue acquisition

Surplus, tumor-adjacent, archived formalin-fixed paraffin-embedded human stomach tissue sections from prior gastrectomies for gastric cancer were acquired from UCSF pathology under IRB 19-27391. Waiver of consent was authorized by the UCSF IRB. Uninvolved, tumor-adjacent gastric tissue at the margin of the resection were selected, and biospecimens were stratified as having received chemotherapy (platin) treatment prior to gastrectomy or not. Patients with *CDH1* mutations were excluded. Tissue sections were not exhausted, and slides were de-identified prior to processing.

#### De-paraffin embed tissue

Slides were washed three times in xylene substitute (Sigma-Aldrich, A5597) for five minutes each wash step. Slides were incubated in 100% ethanol for five minutes, 95% ethanol for five minutes, 70% ethanol for five minutes, and finally deionized water for five minutes.

#### Antigen retrieval

Slides were placed in Antigen Unmasking Solution, Citrate-Based (Vector Laboratories, H-3300-250) in a glass coplin jar. The glass coplin jar was placed in a beaker filled with water and microwaved until the buffer began boiling. The tissue was microwaved for another minute with power set at 50%. The tissue was allowed to cool for a minimum of 30 minutes at room temperature until proceeding with staining.

#### Staining

Tissue was blocked in 1% triton X and 1% BSA for one hour. Tissue was incubated in primary antibody solution diluted in 1% triton X and 1% BSA overnight at 4°C. Tissue was washed three times with 1% triton X and 1% BSA for 30 minutes each wash step. Tissue was incubated in secondary antibody solution diluted in 1% triton X and 1% BSA for 1 hour at room temperature. Tissue was washed three times with 1% triton X and 1% BSA for 30 minutes each wash step. Tissue was mounted in Vectashield HardSet Antifade Mounting Medium (Vector Laboratories, H-1400).

#### Imaging and analysis of p-PKC

High-throughput imaging was carried out using the ImageXpress Confocal HT.ai (Molecular Devices). Neuronal p-PKC levels were detected based on the overlap of p-PKC and TUBB3 signals with the MetaXpress High-Content Image Acquisition and Analysis Software. Representative images were taken with the ImageXpress Confocal HT.ai (Molecular Devices) and processed using Adobe Photoshop software. Neuronal p-PKC integrated intensities were normalized by TUBB3 total area. Each dot represents a region of interest. Data were collected from five regions of interest from eight patients. Statistical analysis was performed by Welch’s t test.

#### Imaging and analysis of nitrergic and cholinergic neurons

High-throughput imaging was carried out using the ImageXpress Confocal HT.ai (Molecular Devices). Nitrergic neurons were detected based on the overlap of NOS1 and HuC/D signals, cholinergic neurons were detected based on the overlap of CHAT and HuC/D signals, and CHAT+ NOS1+ neurons were detected based on the overlap of nitrergic neurons with cholinergic neurons with the MetaXpress High-Content Image Acquisition and Analysis Software. Representative images were taken with the ImageXpress Confocal HT.ai (Molecular Devices) and processed using Adobe Photoshop software. NOS1+ CHAT-neurons were calculated by subtracting the nitrergic neuron total number by the CHAT+ NOS1+ neuron total number. CHAT+ NOS1-neurons were calculated by subtracting the cholinergic neuron total number by the CHAT+ NOS1+ neuron total number. Each of the three populations (NOS1+ CHAT-neurons, CHAT+ NOS1-neurons, and CHAT+ NOS1+ neurons) were normalized by their combined total number to calculate each population’s percent representation. Each dot represents a region of interest. Data were collected from five regions of interest from eight patients. Statistical analysis was performed by Welch’s t test.

### Mouse Treatment Protocols

#### Preparation of solutions for injection

All solutions were prepared fresh for each injection day. A 1 mM stock solution of oxaliplatin (Sigma-Aldrich, Y0000271) was prepared by dissolving it in sterile water by vortexing vigorously and then was sterilized by filtration with a 0.22 µm filter prior to injection. Oxaliplatin’s vehicle was prepared by filtering sterile water with a 0.22 µm filter prior to injection. A 71.1 mg/mL stock solution of oxybutynin (SelleckChem, S1754) was prepared in DMSO, then diluted to 3.6 mg/mL (5%) in 2% tween80 in sterile saline and was sterilized by filtration with a 0.22 µm filter prior to injection. Oxybutynin’s vehicle was prepared by diluting DMSO (5%) in 2% tween80 in sterile saline and was sterilized by filtration with a 0.22 µm filter prior to injection.

#### Intraperitoneal injections

Mice received intraperitoneal injections every three days for five total injection days. Vehicle-treated mice received injections of oxaliplatin’s vehicle (3 mg/kg per dose) and oxybuytnin’s vehicle (35 mg/kg per dose). Oxybutynin-treated mice received injections of oxaliplatin’s vehicle (3 mg/kg per dose) and oxybuytnin (35 mg/kg per dose). Oxaliplatin-treated mice received injections of oxaliplatin (3 mg/kg per dose) and oxybutynin’s vehicle (35 mg/kg per dose). Oxaliplatin-oxybutynin co-treated mice received injections of oxaliplatin (3 mg/kg per dose) and oxybutynin (35 mg/kg per dose). *GI transit time.* Mice were gavaged with 0.2 ml of dye solution containing 6% carmine, 0.5% methylcellulose, and 0.9% NaCl, using a #24 round-tip feeding needle. The needle was held inside the mouse esophagus for a few seconds after gavage to prevent regurgitation. One hour later, the stool color was monitored for gavaged mice every 10 minutes. For each mouse, total GI transit time is between the time of gavage and the time when red stool is observed. Each dot-line pairing represents an individual mouse. Statistical analysis was performed by two-way ANOVA with Šidák correction for multiple comparisons.

### Preparation of Whole Mount Mouse Colon Tissue

Following the excision, the entire colon was pinned in a Sylgard (Dow) lined petri dish and opened along the mesenteric border. The tissue was fixed in 4% paraformaldehyde (Santa Cruz, sc-281692) in PBS for approximately 45 minutes at room temperature. The proximal half of the colon was separated from the distal half, and both were stored in PBS at 4°C.

### Whole Mount Imaging and Analysis

#### Staining

Tissue was blocked in 1% triton X and 1% BSA for one hour with rocking and incubated in primary antibody solution diluted in 1% triton X and 1% BSA overnight at 4°C. Tissue was washed three times with 1% triton X and 1% BSA for 30 minutes each wash step with rocking. Tissue was incubated in secondary antibody solution diluted in 1% triton X and 1% BSA for one hour at room temperature. Tissue was washed three times with 1% triton X and 1% BSA for 30 minutes each wash step with rocking. Tissue was stored in PBS at 4°C.

#### Imaging

High-throughput imaging was carried out using the ImageXpress Confocal HT.ai (Molecular Devices). Nitrergic neurons were detected based on the overlap of NOS1 staining with HuC/D staining, cholinergic neurons were detected based on the overlap of CHAT staining and HuC/D staining, and the total number of neurons was measured by HuC/D staining with the MetaXpress High-Content Image Acquisition and Analysis Software. The proportion of nitrergic neurons was calculated by normalizing the total number of nitrergic neurons by the total number of neurons. The proportion of cholinergic neurons was calculated by normalizing the total number of cholinergic neurons by the total number of neurons. Each dot represents an individual mouse. Statistical analysis was performed by Brown-Forsythe and Welch one-way ANOVA with Dunnett T3 correction for multiple comparisons. Representative images were taken with the ImageXpress Confocal HT.ai (Molecular Devices) and processed using Adobe Photoshop software.

### Generating Figure Schematics

Schematics for the figures were generated using BioRender. https://biorender.com/

